# The genetic consequences of historic climate change on the contemporary population structure of a widespread temperate North American songbird

**DOI:** 10.1101/2024.02.18.580918

**Authors:** Alison Cloutier, David Tsz Chung Chan, Emily Shui Kei Poon, Simon Yung Wa Sin

## Abstract

Studies of widely distributed species can offer insight regarding how past demographic events tied to historic glaciation and ongoing population genetic processes interact to shape contemporaneous patterns of biodiversity at a continental scale. In this study, we used whole-genome resequencing to investigate the current population structure and genetic signatures of past demographic events in the widespread migratory American goldfinch (*Spinus tristis*). In contrast to the low variation in mitochondrial genomes, a genome-wide panel of >4.5 million single nucleotide polymorphisms (SNPs) strongly supported the existence of eastern and western populations separated by western mountain ranges and additional population structuring within the western clade. Demographic modeling indicated that the eastern and western populations diverged approximately one million years ago, and both populations experienced subsequent population bottlenecks during the last glacial period. Species distribution models showed a severe contraction of suitable habitat for the American goldfinch during this period, with predicted discontinuities that are indicative of multiple, isolated glacial refugia that coincide with present-day population structure. This study highlights the power of genome-level sequencing approaches to deepen our understanding of evolutionary processes in nonmodel wild species and to contribute to efforts assessing how historic demographic events and contemporary factors might influence biodiversity.

## Introduction

Understanding how historic demographic events and contemporary population genetic processes contribute to shaping current patterns of biodiversity is receiving renewed focus in view of the threats posed by anthropogenic climate change and landscape-scale alterations to habitat (Cristofari et al. 2018, Gagnaire 2020, McCaslin and Heath 2020). In temperate North America, Quaternary ice ages are recognized as a major historic influence that shaped current species distributions and population structure (Hewitt 2000, 2004). Alternating cycles of glaciation interspersed with periods of interglacial warming during the Pleistocene profoundly reshaped the available habitat for many species and caused populations to contract into refugia located at the southern edges of advancing glaciers before expanding into newly available habitat upon glacial retreat (Hewitt 2000, 2004; Brunsfeld et al. 2001). These historic responses to a changing climate in many cases left tell-tale signatures in the genomes of current species that can be used to infer past demographic events such as population bottlenecks associated with refugial contraction or population expansions that followed glacial retreat (Milà et al. 2000, Spellman et al. 2007, Nadachowska-Brzyska et al. 2015). By investigating these genomic signatures, we can better understand the extent to which current genetic diversity and population structure result from responses to past climatic events and, in turn, we may be better able to predict how current distributional shifts in response to climate change might influence community structure, extinction risk, and wildlife disease dynamics (Cristofari et al. 2018, Mable 2019, Dierickx et al. 2020, McCaslin and Heath 2020, Theodoridis et al. 2020, Smith et al. 2021; Huynh et al. 2023).

However, current genetic diversity is not merely a static byproduct of deep demographic history but is also a dynamic trait that both reflects and influences ongoing processes of migration, genetic drift, and selection at the population level (Wolf et al. 2010, Miraldo et al. 2016; Sin et al. 2021). Studying these processes, at least at the whole-genome level, has traditionally been the purview of comparative population genomics, but the value of phylogeographic methods that interpret patterns of genetic variation in a geographically explicit manner has long been recognized (Avise et al. 1987, Avise 2000, Gagnaire 2020, Edwards et al. 2022). A major focus of comparative phylogeographic approaches is to identify commonalities in the population structure and divergence times of co-distributed taxa to understand how landscape features have shaped current species assemblages (Burbrink et al. 2016, Edwards et al. 2022). Nevertheless, intraspecific studies, especially of species whose ecology, behavior and life history are well characterized, can not only generate baseline data necessary for future comparative studies but can also yield a fine-grained view of the mechanics of selection underlying processes of local adaptation and speciation. Decreasing costs associated with high-throughput sequencing (HTS) have made it possible to adopt genome-level approaches for nonmodel species in the wild and are allowing investigations to expand beyond the study of small numbers of candidate genes to consider how genome architecture itself contributes to a species’ genetic diversity (Wolf et al. 2010, Burri et al. 2015, Manthey et al. 2021, Huynh et al. 2023; Moreira et al. 2023).

Although sometimes overlooked relative to species of conservation concern, common and widespread species can offer insight regarding how demographic events have shaped continental-scale biodiversity, and widespread species often harbor geographically structured phenotypic or behavioral variation whose basis can be investigated at the genomic level. The American goldfinch (*Spinus tristis*, order Passeriformes, family Fringillidae) is a widespread granivorous passerine common to weedy fields, floodplains, and open woodlands across the entire breadth of temperate North America (McGraw and Middleton 2020). With an estimated census size of 44 million breeding individuals (Partners in Flight, 2020), the American goldfinch is a common species throughout much of its range and is categorized as of Least Concern according to the IUCN Red List of Threatened Species (IUCN 2016). *S. tristis* is distributed as far north as central Canada during the breeding season, and as far south as central Mexico (in the east) and northern Baja California (in the west) during the winter, with a wide band stretching across the middle of this distributional range where *S. tristis* occurs year-round (McGraw and Middleton 2020). The American goldfinch is a partial migrant, where some individuals migrate southward to overwinter whereas others remain in sedentary year-round resident populations (Prescott and Middleton 1990). Banding data has shown that even in portions of its range where the American goldfinch occurs year-round, some local breeding populations migrate south in winter and are replaced by distinct overwintering populations (Middleton 1978).

There are four described subspecies of American goldfinch that occupy partially overlapping distributional ranges and that are distinguished by clinal variation in body size and mantle color, in addition to differences in the extent of white or pale wing and tail markings (McGraw and Middleton 2020). *S. t. tristis* (Linnaeus 1758) is the most common subspecies and is found throughout eastern North America as far west as Colorado (Fig. 1). *S. t. pallidus* (Mearns 1890) is distributed across the Great Plains, Rocky Mountains, and Intermountain West. The remaining two subspecies occur along the Pacific coast and coastal mountains, with *S. t. jewetti* (van Rossem 1943) breeding in the Pacific Northwest and overwintering as far south as California and *S. t. salicamans* (Grinnell 1897) largely resident along the Pacific coast from southern Oregon through Southern California, but with some individuals overwintering as far south as northwestern Baja California and as far east as the Mojave and Colorado Deserts (McGraw and Middleton 2020).

**Fig. 1.**
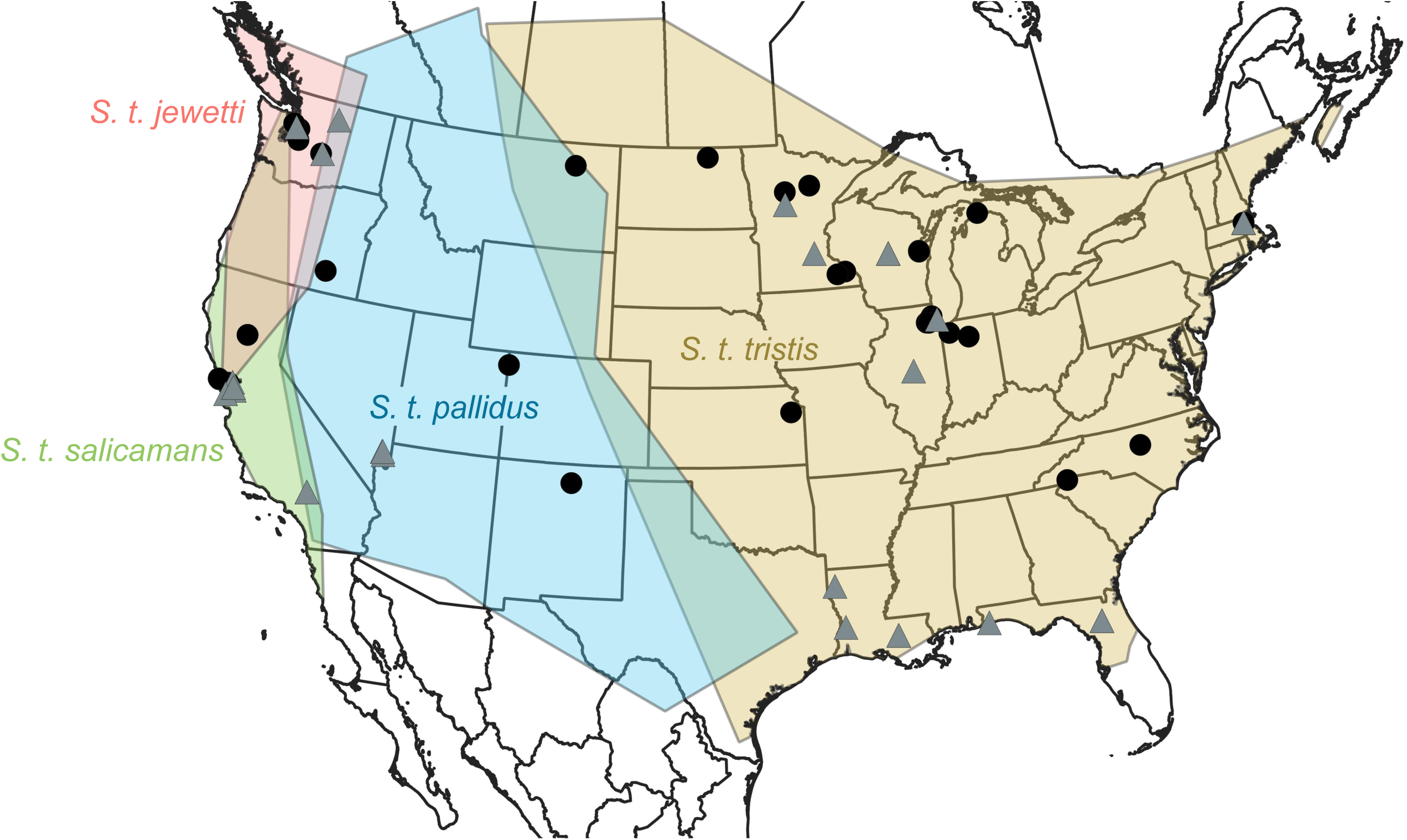
Subspecies range and sampling map for *Spinus tristis*. Shaded polygons indicate distributions of the four described *S. tristis* subspecies estimated from occurrence data obtained from VertNet (Constable et al. 2010). Filled symbols indicate the sampling locations of individuals used for whole-genome resequencing, with black circles indicating individuals sampled during the breeding season and gray triangles indicating individuals sampled during the nonbreeding season.

Despite being a common species and the focus of research spanning topics as varied as cold tolerance (e.g. Cheviron and Swanson 2017), carotenoid-based pigmentation (e.g. Kelly et al. 2012) and the effects of herbicide use (Sughrue et al. 2008), there have been no previous efforts to study the evolutionary history and population differentiation of the American goldfinch with molecular sequence data. In this study, we generate whole-genome resequencing data for 70 individuals sampled across the distributional range of *S. tristis* to investigate population structure using complete mitochondrial DNA (mtDNA) genomes and genome-wide single nucleotide polymorphism (SNP) markers. Our aims were: 1) to assess the extent to which patterns of genetic differentiation coincide with the described *S. tristis* subspecies, 2) to examine how the demographic history of *S. tristis* might have influenced its current genetic variation, and 3) to investigate whether regions of the genome under selection coincide with genes likely to underlie observed phenotypic or migratory differences among populations.

## Materials and Methods

### Whole-genome resequencing

Genomic DNA was isolated from blood or tissue samples from 70 *S. tristis* individuals obtained from museum collections or additional field sampling dating between 1983–2017 (Fig. 1, Suppl. Data 1). Samples were chosen to represent the distributional range of described subspecies as well as to cover both the breeding and nonbreeding periods. DNA was extracted using the E.Z.N.A. Tissue DNA Kit (Omega Bio-tek Inc., Norcross, Georgia USA) following the manufacturer’s protocol and DNA concentration was measured with the Qubit DNA Assay Kit (Thermo Fisher Scientific, Waltham, Massachusetts USA). Libraries of 350 bp insert size were prepared and whole-genome resequencing was conducted by Novogene (Hong Kong). Libraries were sequenced in paired-end mode (2 x 150 bp reads) on an Illumina NovaSeq platform to yield approximately 20 Gb of data per sample. Raw sequencing quality was assessed with FastQC v. 0.11.8 (Andrews 2010).

### Data Analysis

#### Subspecies range mapping

Occurrence data for each of the four described subspecies of *S. tristis* was downloaded from VertNet (Constable et al. 2010, accessed June 30, 2023) and filtered to retain ‘Bird specimens’ (i.e. omitting nest records and acoustic data). Decimal latitude and longitude coordinates were used to build species distribution polygons with the alpha shape hull transformation in the MapMaker application of ModestR v. 6.5 (García-Roselló et al. 2013). The resulting shape files were visualized with QGIS v.

3.32.0 (QGIS Development Team 2021).

#### Read mapping and variant calling

Fastp v. 0.23.2 (Chen et al. 2018) was used to trim adapter sequences and to merge overlapping read pairs. Processed reads were mapped to an existing goldfinch reference genome assembly (S. Y. W. Sin, manuscript in prep.) with BWA-MEM v. 0.7.17 (Li and Durbin 2009). SAMtools v. 1.14 (Li et al. 2009, Danecek et al. 2021) was used to merge and sort mapped reads prior to marking and removing duplicates with Picard v. 2.26.6 (http://broadinstitute.github.io/picard). Mosdepth v. 0.3.4 (Pedersen & Quinlan 2018) was used to calculate mapping depth of coverage (DoC).

Variants were called using Genome Analysis Toolkit (GATK) HaplotypeCaller v. 4.2.4.0 (Van der Auwera and O’Connor 2020) followed by GATK GenomicsDBImport and GenotypeGVCFs modules to output variant call format (VCF) files for individual scaffolds. BCFtools v. 1.14 (Danecek et al. 2021) was used to concatenate output into a single VCF and to filter data to retain SNP variants only and to exclude scaffolds that aligned to sex chromosomes of the zebra finch with SatsumaSynteny (see below). VCFTools v. 0.1.17 (Danecek et al. 2011) was used to filter goldfinch scaffolds below 1 Mb in length and to retain biallelic SNPs only. BCFtools and GATK’s VariantFiltration module were then used to implement the Genome Analysis Toolkit best practices pipeline for variant calling (Van der Auwera et al. 2013, Van der Auwera and O’Connor 2020) to filter SNPs with: QualByDepth (QD) < 2.0, FisherStrand (FS) > 60.0, StrandOddsRatio (SOR) > 3.0, RMS Mapping Quality (MQ) < 40.0, Mapping Quality Rank Sum Test (MQRankSum) < -12.5, and ReadPosRankSum < -8.0 or > 8.0. VCFTools was used to filter SNPs at a minor allele frequency (MAF) threshold of 0.05, maximum missing threshold of 0.95 (i.e. allowing up to 5% missing genotypes per SNP), minDP 3, min-meanDP 3, maxDP 22 and max-meanDP 22 (with the maximum values set at 1.5X the average SNP depth of coverage [DoC] across individuals). PLINK v. 1.90b6.24 (Chang et al. 2015) was used to prune filtered SNPs by linkage disequilibrium with the settings --indep-pairwise 50 10 0.5 (50 Kb window size, 10 bp step size, and r^2^ threshold 0.5). VCFTools was used to output summary metrics for data set quality assessment, including measures of heterozygosity, missingness, and depth of coverage. NGSrelate v. 2 (Hanghøj et al. 2019) was used to calculate pairwise coefficients of relatedness between individuals.

SatsumaSynteny v. 3.0 (Grabherr et al. 2010) was used to align scaffolds of the reference goldfinch genome assembly to the representative RefSeq genome for the zebra finch (*Taeniopygia guttata* assembly bTaeGut1.4pri, GenBank accession GCF_003957565.2). Goldfinch scaffolds aligning to zebra finch sex chromosomes were excluded from further analyses. Remaining goldfinch scaffolds greater than 1 Mb in size were ordered and oriented into autosomal pseudochromosomes according to their alignment with zebra finch chromosome assemblies and zebra finch coordinates were lifted over to the goldfinch for positions that had unique reciprocal alignments between the two genomes.

### Population structure and genetic diversity

Principal components analysis (PCA) of the filtered SNP data set was conducted with PLINK and the results were visualized using the ggplot2 package in R v. 4.1.2 (Wickham 2016, R Core Team 2021). ADMIXTURE v. 1.3.0 (Alexander et al. 2009) was used to infer population structure by assigning individuals to a pre-defined number of genetic clusters (K). Ten replicates were run for each of K=1 through K=6, with five-fold cross-validation and 200 bootstrap replicates. The optimal cluster value was determined based on the lowest cross-validation error as well as using the Best K method of Evanno et al. (2005) implemented in CLUMPAK v. 1.1 (Kopelman et al. 2015). ADMIXTURE clustering results were summarized with CLUMPAK and visualized with the R package pophelper (Francis 2017). VCFTools was used to calculate genome-wide estimates of Tajima’s D in 20 Kb windows and nucleotide diversity (π) in 20 Kb sliding windows with a 10 Kb step size.

### Mitochondrial DNA (mtDNA) assembly and phylogeny

Trimmomatic v. 0.39 (Bolger et al. 2014) was used for adapter removal and read trimming with options ILLUMINACLIP:TruSeq3-PE-2.fa:2:30:10:1:true. 30 million read pairs per individual were subsampled with BBmap v. 39.01 (Bushnell 2014) and were iteratively mapped to the complete mitochondrial genome of the lesser goldfinch (*Spinus psaltria*, GenBank accession NC_025627) with MITObim v. 1.9.1 (Hahn et al. 2013). A consensus *S. tristis* mtDNA genome was computed in Geneious Prime v. 2023 (https://www.geneious.com) from the MITObim assemblies for all individuals and the full set of trimmed reads for each individual was then mapped against this goldfinch consensus reference with BWA-MEM. Mapped reads were processed with SAMtools to retain reads with minimum mapping and base qualities of 30 and duplicates were marked and removed with Picard. A final consensus mtDNA genome was then output for each individual in Geneious, and MAFFT v. 7.520 (Katoh and Standley 2013) was used to align the *S. tristis* mtDNA sequences along with the *S. psaltria* outgroup.

ModelFinder (Kalyaanamoorthy et al. 2017) was used to determine the best-fitting substitution model and partitioning scheme for phylogenetic inference (determined as: partition1 containing the control region [CR], tRNAs, rRNAs, and 1^st^ codon positions of protein-coding genes with a TIM2+F+I substitution model; partition2 with 2^nd^ codon positions and TN+F+I model; partition3 with 3^rd^ codon positions and TPM3+F+G4 substitution model). The maximum-likelihood tree was inferred with IQ-TREE v. 2.1.3 (Minh et al. 2020) from 100 searches beginning with random starting trees, and confidence estimates were obtained from 500 nonparametric bootstrap replicates. Bootstrap values were drawn on the maximum-likelihood tree with RAxML v. 8.2.11 (Stamatakis 2014) and the topology was outgroup rooted with *S. psaltria* using ETE3 (Huerta-Cepas et al. 2016).

### Nuclear DNA phylogeny

BCFtools was used to output individual SNP genotypes from the final filtered VCF and heterozygous sites within an individual were replaced with their IUPAC code. Sequence for the outgroup zebra finch was compiled using coordinates lifted over from the SatsumaSynteny whole-genome alignment detailed above.

The maximum-likelihood topology was inferred in IQ-TREE from a starting set of 100 trees and a general time reversible (GTR) substitution model. Confidence was assessed with 100 nonparametric bootstrap replicates. Bootstrap replicates were plotted on the maximum-likelihood tree with RAxML and the topology was outgroup rooted with the zebra finch using ETE3.

In addition to the ‘base’ data alignment matrix detailed above, we also tested the effect of replacing IUPAC codes with ‘called’ bases for sites where the most frequent allele within an individual occurred at > 2X that of the less common allele, as well as using trimAl v. 1.2rev59 (Capella-Gutiérrez et al. 2009) to trim alignment columns that consisted of >10%, >20%, or >30% missing data.

### Demographic history

The pairwise sequential Markovian coalescent (PSMC) was used to reconstruct the demographic histories of the eastern and western goldfinch populations. Sequencing reads from the individual used for the reference *de novo* whole-genome assembly (*S. t. tristis*, S. Y. W. Sin, manuscript in prep.), with an average 48X genomic depth of coverage, served as the representative of the eastern population. GOFI_59, which was sampled from Washington state (Suppl. Data 1), was used as the western population representative. A second 350 bp insert sequencing library (138,048,084 read pairs, GenBank accession [pending]) was constructed and sequenced in paired-end mode in a partial Illumina NovaSeq lane to increase genome average depth of coverage for this sample to 35X.

Sequencing reads were pre-processed with fastp and mapped to the reference goldfinch assembly with BWA-MEM as described above. SAMtools was used to filter reads with mapping quality < 30 and Picard was used to mark and remove duplicates. SAMtools was then used to randomly subsample mapped reads for the eastern individual to reduce average DoC to 35X to match that of the western individual, and to retain only autosomal scaffolds greater than 1 Mb in length for both individuals. Consensus calling was done with BCFtools mpileup with minimum base and mapping qualities of 30, followed by BCFtools consensus calling (‘call -c’ option), and the vcf2fq utility with minimum base quality 30 and sequencing depth thresholds set according to the PSMC developer’s recommendations (https://github.com/lh3/psmc, minimum depth of 1/3 the average [option -d 11] and maximum depth 2X the average [option -D 70]).

PSMC v. 2016.1.21 (Li and Durbin 2011) analyses were run with an atomic time interval parameter found to be appropriate for a wide range of avian species (Nadachowska-Brzyska et. al 2015, parameter -p 4+30*2+4+6+10) and confidence was assessed with 100 bootstrap replicates.

EasySFS (https://github.com/isaacovercast/easySFS, Gutenkunst et al. 2009) was used to compute the folded site frequency spectrum (SFS) for use in two additional methods of demographic inference, Stairway Plot 2 and fastsimcoal2. Site frequency spectra were generated from SNP data with no minor allele frequency (MAF) filtering applied and values were projected down to 86 haplotypes (of 96 total) for the eastern population and 34 haplotypes (of 38 total) for the western population to maximize the total number of segregating sites while retaining a sufficiently large sample size for analysis (Gutenkunst et al. 2009).

Stairway Plot 2 v. 2.1.1 (Liu and Fu 2015, 2020) was run separately for the eastern and western goldfinch populations using the two-epoch model with the default 67% of sites used for training, 200 input files created for estimation of pseudo-CIs, and the default four values for the number of random break points for each trial as specified by the authors.

A second SFS-based method, the fast sequential Markov coalescent (fastsimcoal2 v. 2702, Excoffier et al. 2013, 2021) was used to estimate demographic parameters under eight simulated models: 1) no migration between eastern and western goldfinch populations, 2) migration in two different time periods, 3) early migration, 4) ongoing migration, 5) recent migration, 6) early migration followed by independent bottleneck events in each population, 7) independent bottlenecks in each population with no migration, 8) independent bottlenecks in each population followed by migration between populations. Each model was run 50 times with 500,000 coalescent simulations and SFS entries with fewer than 10 SNPs were pooled. The run with the highest estimated likelihood was chosen for each candidate model, and candidate models were compared according to the Akaike information criterion (AIC). Confidence intervals for estimated parameters in the best candidate model were obtained from 10 parametric bootstrap resamplings of the input SFS data.

Results of all demographic analyses were scaled with the estimated mutation rate for the medium ground finch (*Geospiza fortis*, 3.4412 x 10^-9^, Nadachowska-Brzyska et al. 2015) since no estimates are available for the American goldfinch. A generation time equal to twice the average age of sexual maturity of the American goldfinch according to the Animal Ageing and Longevity Database was used (Tacutu et al. 2018, generation time = 2 years).

### Species distribution modeling

Species distribution models (SDMs) were generated using MaxENT v.3.4.4 (Phillips et al. 2017) to investigate the potential effect of paleoclimate on the distribution of *S. tristis*. Occurrence data for *S. tristis* were downloaded from eBird (eBird 2021). Only data from 1979 to 2013 were retained to match available climate data from PaleoClim (Brown et al. 2018), and occurrence data for nonbreeding (January to February) and breeding (July to August) periods were analyzed separately. Occurrence data were further filtered to retain observations from the middle of each breeding and nonbreeding season to account for individual variation in migration time and to ensure individuals were correctly assigned to their breeding or nonbreeding grounds. Data lacking GPS coordinates were removed and filtered data were rarefied to avoid overfitting through removal of samples located within 40 km of each other with SDMtoolbox (Brown et al. 2017). A total of 26,409 points for breeding season and 22,564 points for nonbreeding season were retained.

Nineteen terrestrial bioclimatic variables were obtained from the PaleoClim database (Brown et al. 2018) for the present time period (1979–2013), Meghalayan (4.2–0.3 thousand years before present, Kbp), Northgrippian (8.326–4.2 Kbp), Greenlandian (11.7–8.326 Kbp), Younger Dryas Stadial (12.9–11.7 Kbp), Bølling-Allerød (14.7–12.9 Kbp), Heinrich Stadial 1 (17.0–14.7 Kbp), Last Glacial Maximum (*c.* 21 Kbp), Last Interglacial (*c.* 130 Kbp), and Marine isotope stage (MIS) 19 (*c.* 787 Kbp). Bioclimatic data were separated into breeding and nonbreeding season based on their description, including climate data of wettest, driest, warmest, and coldest months or quarters since there are no paleoclimatic data per month available from PaleoClim. Global climate data were cropped to the North American continent (15°03’33.6”N to 71°21’55.9”N and 53°59’18.2”W to 164°51’26.3”W). Bioclimatic parameters with a correlation coefficient larger than 0.8 were removed before inferring SDMs for all environmental factors for nonbreeding and breeding seasons, and factors were removed based on jackknife, factor ROC curve, and model AUC. Predictors Bio 1, Bio 8, Bio 12, and Bio 15 were retained for the breeding season, and Bio 1, Bio 8, Bio 15, and Bio 19 were retained for the nonbreeding season. The final SDMs were generated using MaxENT with 50 replicates for each model. Model output and spatial environmental data were visualized in QGIS (QGIS Development Team 2021).

### Fst outlier analysis

We compared per-site Fst values between the eastern and western goldfinch populations and between the two subclades we identified within the western population (see Fig. 3). OutFLANK v. 0.2 (https://github.com/whitlock/OutFLANK) was used to calculate per-site Fst and to identify statistical outliers by fitting a Chi-square distribution to the data and applying a multiple test correction at a q-value threshold of 0.05. BEDTools v. 2.31.0 (Quinlan and Hall 2010) was used to identify annotated genes lying within 1 Kb of outlier SNPs. Statistical overrepresentation of Gene Ontology (GO) terms for biological process, molecular function, and cellular component among outlier SNPs was assessed with the PANTHER classification system v. 18.0 (Mi et al. 2019, Thomas et al. 2022) employing Fisher’s exact tests and false discovery rate (FDR) correction.

## Results

### Whole-genome resequencing and variant calling

Approximately 70 million read pairs were obtained for each of the 70 *S. tristis* whole-genome resequencing libraries (Suppl. Data 2, mean= 71.3, range= 63.9–77.3 million read pairs per library), representing on average 11.8X genomic depth of coverage (DoC) when mapped to the reference genome assembly (Suppl. Data 2, range=8.7–14.0X DoC excluding omitted samples, see below).

No close relatives were identified among sequenced individuals (all pairwise relatedness values < 0.035). However, an initial run-through of the variant calling pipeline and principal components analysis (PCA) identified three outlier individuals: GOFI_07, GOFI_23, and GOFI_57. These samples had the highest proportion of missing SNPs (>25% of sites) and the lowest average SNP depth of coverage (<5.5X DoC) and were therefore excluded from the data set before re-running the final variant calling pipeline and performing all downstream analyses.

The final SNP dataset comprised 4,783,622 biallelic SNPs with an average 96% of SNPs called per individual (range= 78.8–98.7%) and average SNP depth of coverage within an individual of 11.2X (range= 5.4–14.4X, Suppl. Data 2).

### Population structure

Principal components analysis (PCA) did not cluster samples according to sampling season or described *S. tristis* subspecies, but instead clearly divided individuals into an eastern group that contained all individuals sampled throughout the *S. t. tristis* and *S. t. pallidus* distributional ranges and a western group that encompassed all individuals from the *S. t. jewetti* and *S. t. salicamans* ranges (Fig. 2a,b). The second principal component axis (PC2) further subdivided the western group into two clusters whereas the eastern group lacked a well-defined second cluster and instead separated out five individuals in a diffuse grouping (comprising individuals GOFI_36, 37, 49, 61, and 62).

**Fig. 2.**
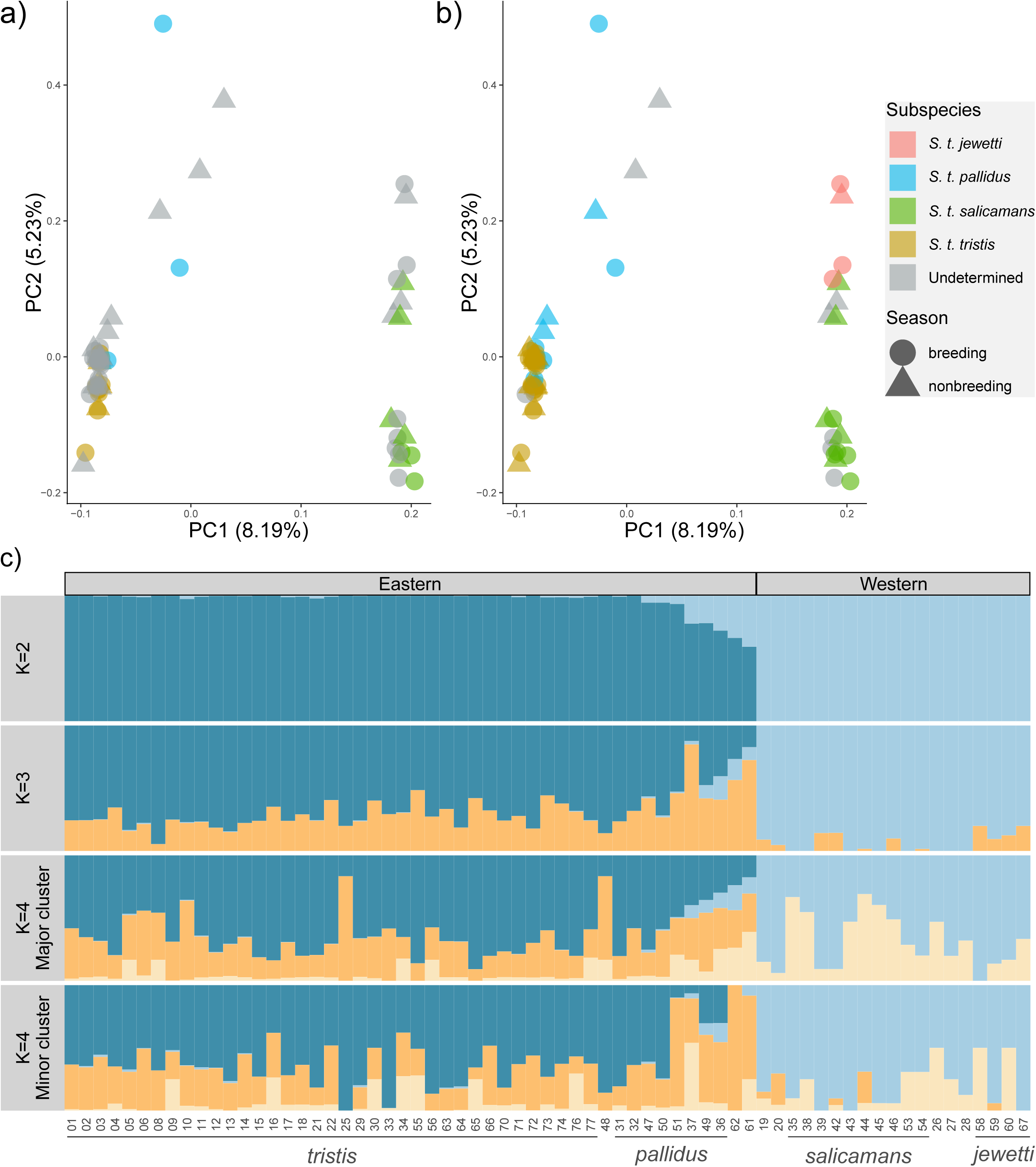
Principal components analysis (PCA; a,b) and ADMIXTURE (c) analysis of population structure in *S. tristis*. Specimens that were identified to subspecies level in museum accession metadata are colored in part a, and those that were identified to subspecies level or that fall clearly within a predicted subspecies distributional range are colored in part b. c) ADMIXTURE results for cluster sizes K=2 to K=4. Replicates for K=4 failed to converge on a single cluster value, and results are shown for major (N= 6 replicates) and minor (N=4 replicates) cluster assignments. The numerical portion of sample IDs are shown along the X-axis. Subspecies designations are shown beneath the x-axis for individuals that were identified to subspecies level or that clearly fell within a predicted subspecies range (corresponding to assignments shown in part b).

The bestK method of Evanno et al. (2005) supported K=2 as the optimal cluster value for ADMIXTURE analyses (Fig.2c), although examination of raw likelihood scores would instead indicate that a single population (K=1) provided the best fit to the data. The K=2 cluster divided individuals into the same eastern and western groups as were observed from PCA axis PC1. Higher values of K failed to distinguish additional groupings identified from principal component PC2, and replicates with K=4 clusters and above failed to converge upon a single cluster assignment. The five individuals listed above that formed a ‘diffuse’ second PCA group for the eastern samples were clearly assigned to the eastern ADMIXTURE cluster at K=2; however, these samples displayed the greatest signatures of genomic admixture.

There were eight individuals with no available subspecies metadata and where assignment to a subspecies range was uncertain due to their presence within or very near to a region of predicted distributional overlap between subspecies (specimens designated ‘undetermined’ in Fig. 2b and not assigned to subspecies in Fig. 2c x-axis labels). Of these, GOFI_48, which was sampled from a region of *S. t. tristis/S. t. pallidus* overlap, and GOFI_26, 27, and 28, which were all sampled near the *S. t. jewetti/S. t. salicamans* boundary, can only be classified to the level of eastern and western populations, respectively. GOFI_19 and GOFI_20, sampled during the nonbreeding period from an area of *S. t. salicamans/S. t. pallidus* overlap, clearly fall within the western group that includes *S. t. salicamans*. GOFI_61 and GOFI_62, both sampled during the nonbreeding period from an area of *S. t. jewetti/S. t. pallidus* overlap are assigned to the eastern population that includes *S. t. pallidus* although, as mentioned above, the cluster assignment proportions for these individuals are the lowest observed.

### Mitochondrial genome assembly and phylogeny

Complete mitochondrial (mtDNA) genomes of 16,814–16,817 bp in length were assembled for all individuals, with all tissue-derived DNA libraries having average depth of coverage >500X and DNA libraries prepared from blood with DoC >16X (Suppl. Data 2). There were 315 variable sites in 16,818 bp of aligned sequence for the ingroup goldfinch specimens (1.9% of total), with only 0.8% of sites being parsimony informative (134 of 16,818 alignment columns).

There was a weak trend for western samples to cluster, with 11 of the 19 individuals from the western group identified from PCA and ADMIXTURE forming a clade with 92% bootstrap support (Fig. S1). However, the mtDNA phylogeny is overall very poorly resolved and is characterized by low bootstrap support across most of the deeper internal branches. Correspondingly, there is no robust support from mtDNA for individuals clustering according to sampling season, described subspecies, or the eastern and western populations identified from PCA and ADMIXTURE analyses.

### Nuclear SNP phylogeny

In contrast to the mtDNA phylogeny, phylogenetic inference using nuclear SNP data recovered a western clade with 100% bootstrap support that was identical to the western population from PCA and ADMIXTURE analyses (Fig. 3a). This western group was further divided into two subclades, each with 100% support, that were identical to the two western clusters delineated by PCA axis PC2 (Fig. 3a). Additional data sets that assessed the effect of alignment filtering or thresholds for adopting IUPAC degenerate codes produced topologies that were identical in recovering this western clade and two subclades with maximum bootstrap support (results not shown).

**Fig. 3.**
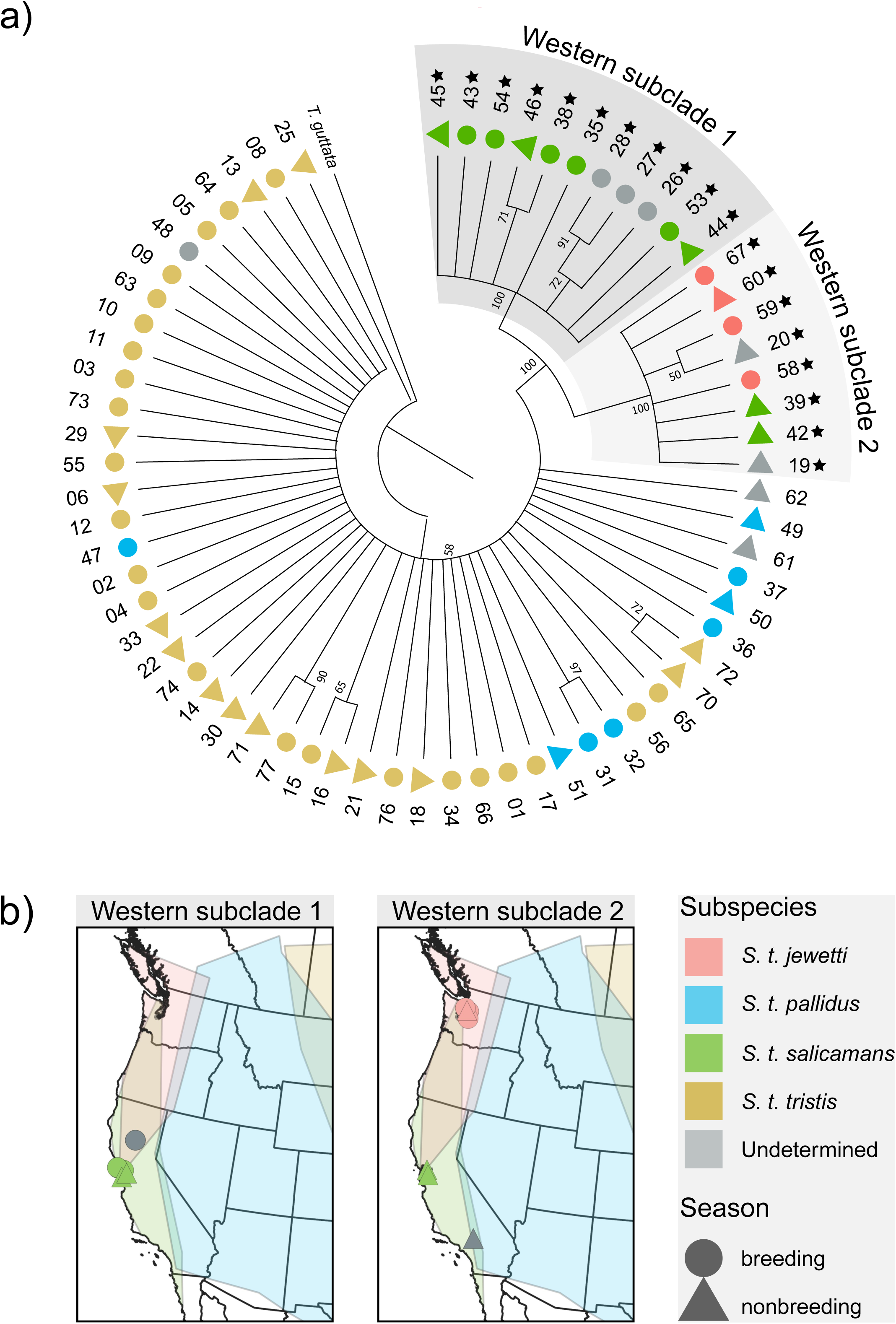
Nuclear SNP phylogeny of *S. tristis* and *Taeniopygia guttata* (zebra finch) outgroup. a) Maximum likelihood phylogeny inferred from SNP data with IQ-TREE, with support from 100 bootstrap replicates indicated. Branch lengths are uninformative, and branches receiving <50% support are collapsed. The numerical portion of sample IDs from Suppl. Data 1 are indicated for each individual. Individuals belonging to the western population from PCA and ADMIXTURE analyses (Fig. 2) are indicated with a star symbol, and the two western subclades found at 100% bootstrap support are labeled and shaded. b) Sampling locations for individuals belonging to the two identified western subclades in part a).

Western subclade 1 (Fig. 3b) contains individuals sampled during both the breeding and nonbreeding seasons along the mid-California Pacific coast (with 6 of these 8 individuals identified as *S. t. salicamans* in museum accession metadata), as well as three individuals sampled during the breeding season from a region of predicted *S. t. salicamans/S. t. jewetti* overlap lying to the northeast (none of these samples was identified to subspecies level). Western subclade 2 (Fig. 3c) contains both breeding and nonbreeding season samples from Washington state (none was identified to subspecies level), as well as two individuals sampled from roughly the same mid-California coastal locality as subclade 1 in the nonbreeding season (and recorded as *S. t. salicamans* in accession metadata) and two individuals sampled from a predicted region of *S. t. salicamans/S. t. pallidus* overlap in Southern California during the nonbreeding season (neither was identified to subspecies level).

Unlike the strongly differentiated western specimens, individuals belonging to the eastern population defined by PCA and ADMIXTURE failed to form a well-supported monophyletic clade in the SNP phylogeny. There was also little evidence for lower-level phylogenetic relationships within this larger eastern sample, with almost all internal branches receiving below 50% bootstrap support (Fig. 3a).

### Demographic analyses

PSMC analysis of single eastern and western representative genomes and Stairway Plot 2 analysis of site frequency spectra (SFS) derived from all sampled individuals yielded similar overall shapes in the estimated demographic histories of each population through time (Fig. 4a,b). PSMC, which is better at inferring more ancient demographic processes (Li and Durbin 2011, Beichman et al. 2018), indicates that the eastern and western populations shared a similar demographic history prior to 1 million years before present (Ybp, Fig. 4a). After this point, the trajectories of the two populations diverge, with the eastern population size increasing more sharply and exhibiting a dip in effective population size (Ne) at the beginning of the last glacial period (LGP) before rebounding to an even higher Ne (*c.* 4 million individuals) that holds steady throughout the remainder of the LGP. The western population exhibits a slower rise in effective population size, peaking at a lower maximum estimate and beginning to decline again prior to the onset of the LGP.

**Fig. 4.**
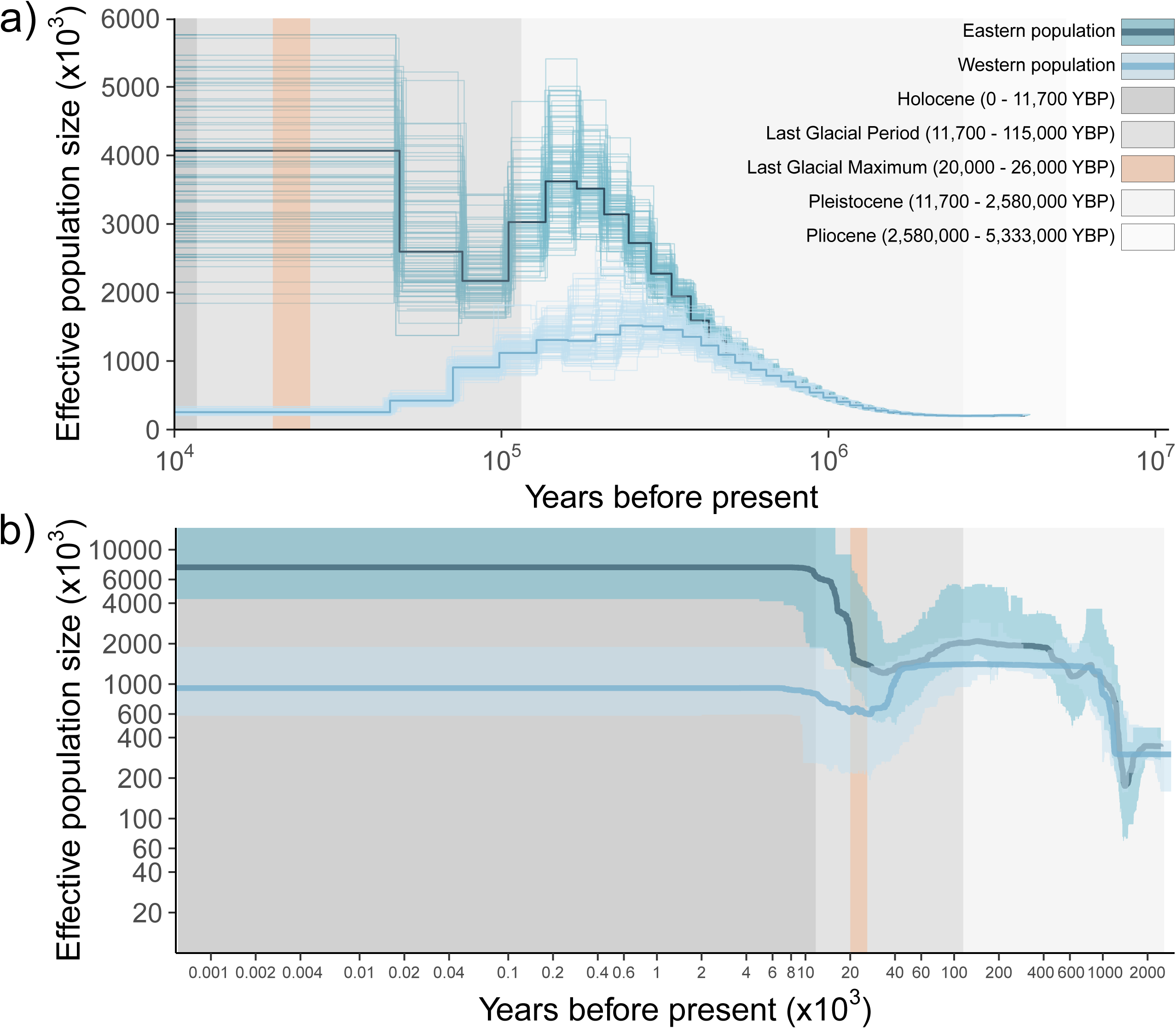
Inference of the demographic history of *S. tristis*. a) PSMC analysis of the genome from a representative individual from the eastern and western populations. b) Stairway Plot 2 analysis of site frequency spectra (SFSs) constructed from individuals used for whole-genome resequencing. Plotted values (light and dark blue lines) show estimated changes in effective population size (Ne) through time, with lighter lines (a) or shaded areas (b) indicating the 95% confidence interval of Ne estimates.

Stairway Plot 2, which is better able to reconstruct more recent population size changes (Liu and Fu 2015, 2020; Beichman et al. 2018), is relatively consistent with PSMC throughout except that both eastern and western populations show a pronounced dip in Ne that coincides with the last glacial maximum (LGM) at 20,000–26,000 Ybp rather than declining at the start of the last glacial period (Fig. 4b). Effective sizes of both populations rebound shortly after the LGM, although predicted Ne of the western population remains below what was predicted prior to the LGM, and estimated sizes of both populations remain stable throughout the remaining time interval to the present (Fig. 4b).

In contrast to these results, fastsimcoal2 identified ongoing migration following population divergence as the best candidate model in preference to models incorporating bottleneck episodes (Table S1, Fig. 5a). Despite this difference, the best fastsimcoal2 model estimated current effective population sizes (*c.* 6.86 million and 0.94 million diploid individuals for the eastern and western populations, respectively) and time to population divergence (*c.* 1 million Ybp) that coincided with the results of PSMC and Stairway Plot 2. Although the best candidate model provides a reasonably good fit overall (Fig. 5b,c), there are some notable areas of divergence between the observed and expected SFS (Fig. 5d), and 95% confidence intervals for both estimated migration rates encompass zero. We attempted to model scenarios that would more closely match the results of PSMC and Stairway Plot 2 to determine if they would provide an improved fit to the observed SFS (for example, modeling independent population expansions after divergence followed by independent bottlenecks and/or population declines). However, most replicates failed to reach convergence for these more parameter-rich models (results not shown).

**Fig. 5.**
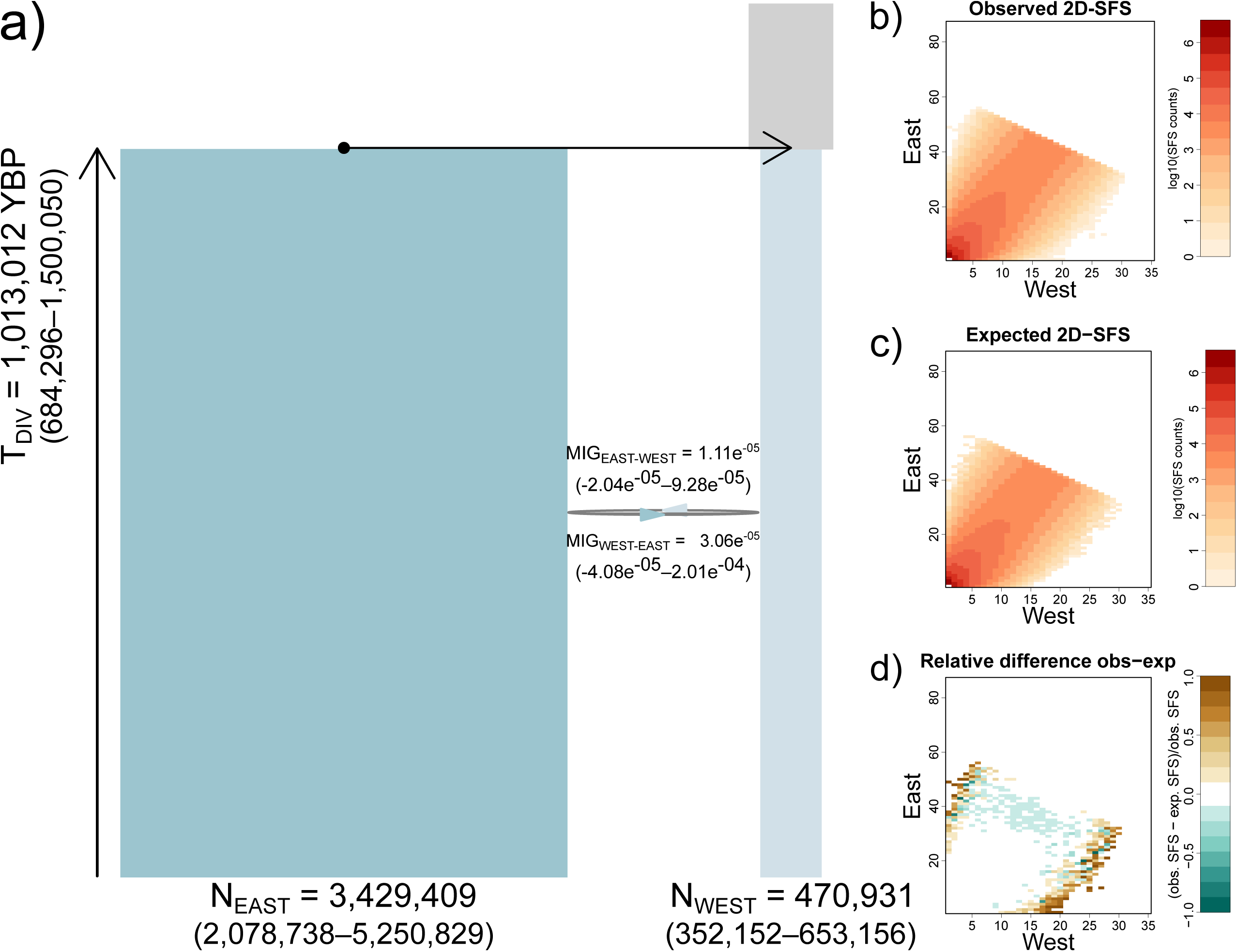
Parameters of the best-fitting model estimated with fastsimcoal2, involving ongoing migration following divergence of the eastern and western populations of *S. tristis*. a) Estimated parameters of the best-fitting candidate model, with 95% confidence intervals shown in brackets beneath each estimate. Migration events are indicated with reciprocal arrows and represent ongoing events following population divergence. N_EAST_/N_WEST_= current effective population sizes of the eastern and western populations, MIG_EAST-WEST_/MIG_WEST-EAST_= migration rate from the eastern to western, or western to eastern population, respectively, T_DIV_= time of ancestral population division into the eastern and western populations. Effective population sizes are given in haploid numbers, and the divergence time is shown as years before present (Ybp). b-d) Assessment of model fit for the best candidate model shown in part a), showing matrix representations of the observed 2D site frequency spectrum (SFS, b), the 2D SFS expected if the candidate model provided a perfect fit to the data (c), and the relative difference between observed and expected SFS matrices (d).

### Species distribution modeling

Current habitat suitability predicted from species distribution modeling largely coincides with the described present range of *S. tristis*, except for a more northerly breeding distribution predicted to extend up the Pacific coast and into Alaska and a more southerly breeding range extending into Mexico, albeit both are associated with low probabilities of occurrence (Fig. 6a). Hindcasts of historical habitat suitability predict contractions to breeding and overwintering distributions during cold glacial periods (e.g. Fig. 6b,c) and expansions during warmer interglacial periods (e.g. Fig. 6d,e). The last glacial maximum (LGM, Fig. 6c) is associated with the most severe constriction to predicted habitat suitability, with suitable breeding and overwintering ranges largely restricted to the Pacific coast and extreme southern United States into Mexico. There is a predicted discontinuity in suitable breeding habitat for *S. tristis* between the eastern and western portions of its range during the LGM, as well as a discontinuity between more northern and southern portions of the breeding range along the Pacific coast (Fig. 6c).

**Fig. 6.**
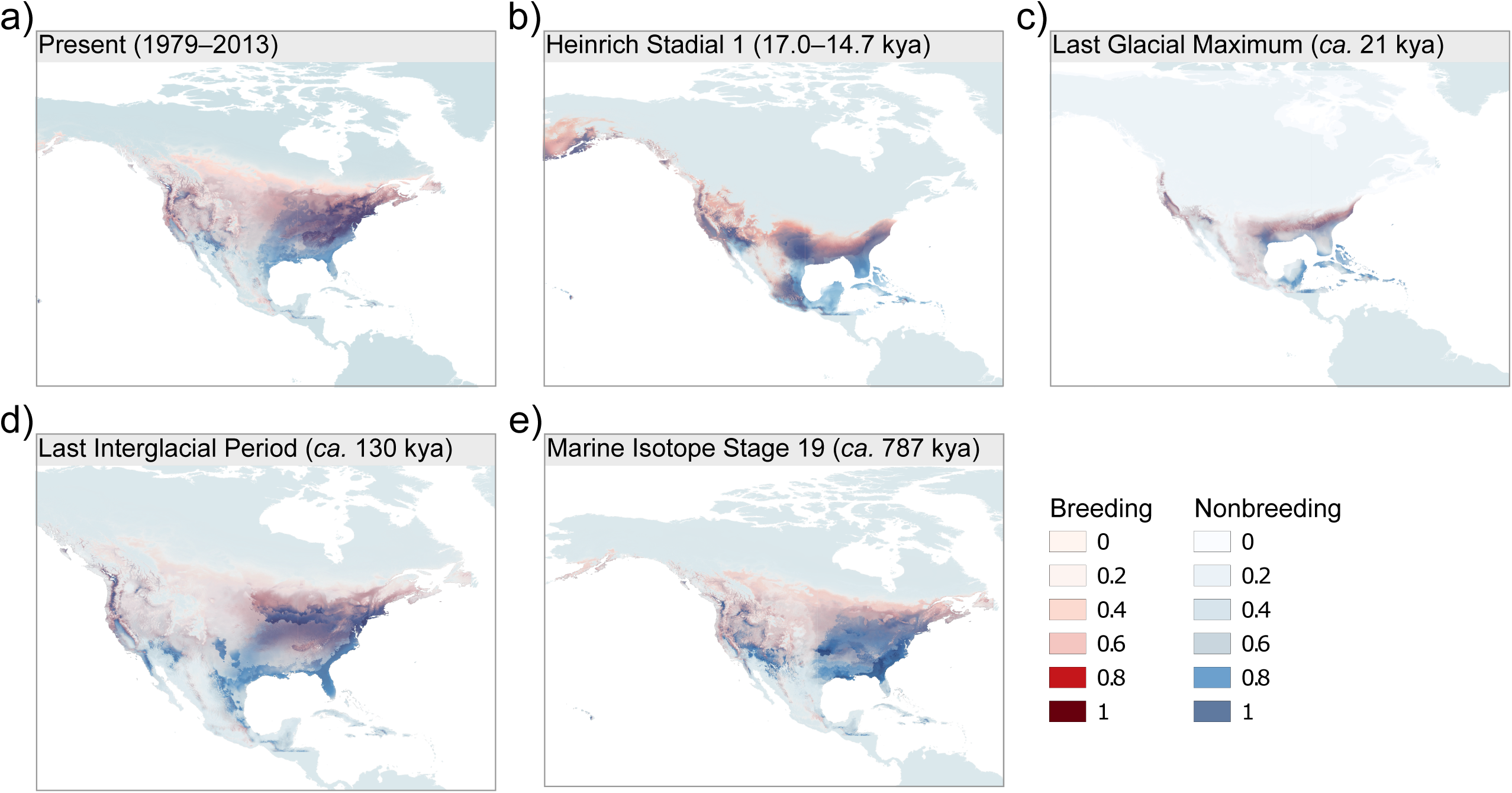
Species distribution models inferred with MaxENT, showing predicted habitat suitability for *S. tristis* during the present time (a) and historical time periods (b-e). Darker colors indicate higher probabilities of species occurrence for a given geographic area.

### Genomic variation and Fst outlier analysis

Mean genome-wide estimates of nucleotide diversity (π) were identical for the eastern and western populations (0.0011 ± 0.0008 standard deviation (SD) for both, Fig. S2a). Mean estimates of Tajima’s D were 0.9254 ± 0.4563 SD for the eastern population and 0.4196 ± 0.4114 SD for the western population (Fig. S2b).

Mean per-site Fst was weak to moderate between the eastern and western populations (0.0132 ± 0.0440 SD, Fig. S3) and very low between the two western subpopulations (0.0026 ± 0.0783 SD, not shown). There were no perfectly segregating SNPs for either the east/west or western subclades comparisons, and there were also no outlier SNPs identified between the western subclades. However, OutFLANK analysis identified 631 outlier SNPs between the eastern and western populations, with 383 of the 631 identified outliers lying within 1 Kb of an annotated gene (291 genes total; some genes contain multiple outlier SNPs, Suppl. Data 3). Statistical overrepresentation tests in PantherDB found no overrepresented Gene Ontology (GO) categories for biological process, molecular function, or cellular component. Closer examination of outlier SNPs falling within annotated genes found that only four SNPs fell within coding regions (one each in DSCAM, FAM199X, HKDC1, and AHSG) whereas the remaining outliers localized to intronic or UTR sequence. Of the four coding SNPs, only one (in AHSG) produced a nonsynonymous substitution, resulting in a semi-conservative replacement of serine by proline.

## Discussion

Although several previous studies used mitochondrial DNA (mtDNA) to investigate phylogenetic relationships and speciation dynamics among goldfinches and siskins (e.g. Arnaiz-Villena et al. 2009, 2012; Beckman and Witt 2015), to the best of our knowledge this study represents the first use of molecular sequencing data to infer the genetic structure and demographic history within the American goldfinch. In addition to a large panel of more than 4.5 million nuclear SNPs, our whole-genome resequencing approach produced a complete mtDNA genome for each sampled individual. We found low intraspecific sequence variation of mtDNA (only 1.9% of sites were variable across the entire genome and only 0.8% were parsimony informative) that provided little support for phylogenetic relationships within *S. tristis*. mtDNA was often the marker of choice in past avian phylogeographic studies and provided resolution of population structure at broad geographic scales (reviewed in Avise and Walker 1998). However, it is suggested that mtDNA variation is likely insufficient to resolve relationships at a finer geographic scale for migratory passerines (Lovette et al. 2004), an assessment that would seem to apply here.

In contrast, we find robust support for intraspecific relationships in *S. tristis* from nuclear sequence data. Our analysis of genome-wide SNP data does not fully support the division of *S. tristis* into its four previously delineated subspecies. Instead, we find evidence for an eastern population that encompasses all individuals sampled from the *S. t. tristis* and *S. t. pallidus* distributional ranges and a western population that includes all individuals sampled from the *S. t. jewetti* and *S. t. salicamans* ranges. The western population extends from central Washington state, Oregon, and east/central California to the Pacific coast, whereas the eastern population covers the entire continental region east of this divide. This pattern strongly suggests that the Cascade and Sierra Nevada Mountain ranges, which extend in a north-south orientation through central Washington and Oregon and east/central California, respectively, represent a geographic barrier between the observed populations. Similar east/west patterns of population differentiation are described for many widespread migratory bird species in temperate North America, with distinct populations often separated by western mountain ranges and/or the Great Plains (e.g. Milot et al. 2000, Kimura et al. 2002, Reugg and Smith 2002, Klicka et al. 2011, Lovette et al. 2004, Aguillon et al. 2018). Together with evidence that historic reductions in effective population sizes of many avian species coincided with the onset of the last glacial period (Nadachowska-Brzyska et al. 2015), this east/west phylogeographic split is proposed to represent genetic differentiation arising from contraction into isolated refugia to either side of a geographic divide during periods of glacial advance (Avise and Walker 1998, Hewitt 2000, Lovette 2005). Our demographic and species distribution modeling support this hypothesis for the American goldfinch, although the timing of inferred events varies somewhat among methods. All three methods of demographic inference date population divergence to the mid-Pleistocene (*c.* 1 million Ybp), following which PSMC analyses infer reductions to effective population size through the start of the last glacial period and Stairway Plot 2 analyses infer population bottlenecks in both the eastern and western populations concurrent with the last glacial maximum (LGM). Species distribution models predict a contraction in suitable goldfinch breeding habitat with an east/west discontinuity compatible with isolated glacial refugia during this period, lending further support that historic climatic conditions likely contributed to shaping current population structure. The moderate nucleotide diversity observed for both the eastern and western populations (π= 0.0011) despite large long-term effective population sizes might in part reflect these historic bottlenecks, although similar estimates of nucleotide diversity are reported for other widespread avian species with large census sizes (e.g. hooded crow, *Corvus (corone) cornix*, π= 0.0011, Dutoit et al. 2017; turkey vulture, *Cathartes aura,* π= 0.0012, Li et al. 2014, Robinson et al. 2016; great black cormorant, *Phalacrocorax carbo,* π= 0.0014, Li et al. 2014, Robinson et al. 2016).

Despite the clear division into eastern and western populations from PCA, ADMIXTURE, and nuclear phylogenetic analyses, five individuals assigned to the eastern population showed greater admixture proportions than the remaining eastern specimens. Aside from one individual sampled in Nevada during the nonbreeding season, these individuals with more admixed genomic signatures were all sampled from locales within, or immediately adjacent to, regions of predicted overlap between eastern and western subspecies (*S. t. pallidus* and *S. t. jewetti*, respectively). Contact zones between the Rocky and Sierra Nevada/Cascade mountains are described for eastern and western populations of several avian taxa (e.g. Barrowclough et al. 2004, Spellman et al. 2007, Manthey et al. 2011), and our results suggest that introgression could occur along at least some portion of this east/west population divide in the American goldfinch. This introgression and/or migration across subspecies contact zones could explain why fastsimcoal2 identified ongoing migration as the best candidate demographic model in preference to models that incorporated bottleneck events. Both PSMC and Stairway Plot 2 inferred bottlenecks of relatively small amplitude and short duration, and mean genome-wide estimates of Tajima’s D were only weakly positive, indicating that more recent migration events might be dominating the genomic signal modeled by fastsimcoal2 coalescent simulations (Yang 2014, Burbrink et al. 2016).

Analysis of genome-wide SNPs identified no finer-scaled structuring within the eastern population of American goldfinches, and we cannot distinguish between the two eastern subspecies, *S. t. tristis* and *S. t. pallidus*, although we note there was relatively sparse sampling across the *S. t. pallidus* range. Earlier studies of widespread North American passerines employing mtDNA markers similarly found low overall genetic diversity and an absence of genetic structuring among sampling sites in eastern continental North America (Kimura et al. 2002, Reugg and Smith 2002, Lovette at al. 2004, Klicka et al. 2011). This absence of genetic differentiation across a broad geographic region could be explained by the loss of variation during bottleneck episodes when populations contract into glacial refugia followed by rapid postglacial expansion (Zink 1996, Hewitt 2000). Weak population structuring, ‘star-shaped’ mtDNA haplotype networks, and latitudinal gradients in intraspecific genetic variation consistent with post-glacial leading-edge expansion all support a role for historic demographic processes linked to climatic cycles in shaping the current patterns of genetic diversity observed for avian taxa across eastern North America (Zink 1996, Hewitt 2000, Lovette et al. 2004, Klicka et al. 2011, Smith et al. 2017). Gene flow between eastern populations might also be acting to limit genetic differentiation across the east. Biogeographic studies of unglaciated eastern North America identified eight phylogeographic discontinuities that could pose a barrier to gene flow. However, common discontinuities were rarely identified for birds or mammals, suggesting that gene flow across these barriers could be sufficient to prevent genetic structuring in highly vagile species such as migratory birds (Soltis et al. 2006, Lyman and Edwards 2022).

Despite this reported absence of genetic structuring in eastern North American migratory passerines, recent investigations incorporating large numbers of nuclear loci are providing a more nuanced view of breeding population structure in some cases. For instance, Colbeck et al. (2008) distinguished an Atlantic breeding population of the American redstart (*Setophaga ruticilla*) that differed from the remaining mainland continental samples, but neither mtDNA nor amplified fragment length polymorphism (AFLP) markers provided robust support for additional structuring within the mainland. More recently, DeSaix et al. (2023a) identified five distinct but weakly differentiated clusters across the *S. ruticilla* breeding range (including the previously described Atlantic/mainland split) by combining dense population sampling with low-coverage whole-genome sequencing. Our whole-genome resequencing approach yielded ample data in terms of the number of SNP markers generated. However, it is likely that additional sampling within the eastern continental region, particularly throughout the Great Plains and American West, is required to definitively assess whether finer-scale structuring exists within the eastern population of the American goldfinch and, if so, to what extent it coincides with the described eastern subspecies.

In contrast to the lack of genetic differentiation in the eastern population, principal components analysis and the nuclear SNP phylogeny, but not ADMIXTURE analysis, clearly discriminate between two western subclades in the American goldfinch. These groupings might correspond to the described western subspecies, as western subclade 2 includes individuals sampled from the *S. t. jewetti* range in Washington state during the breeding period and subclade 1 contains both breeding and nonbreeding season individuals sampled from California, where *S. t. salicamans* typically occurs as a year-round resident. Additional sampling across the western distribution of *S. tristis* is needed to investigate the full extent of spatial structuring across this region, and to determine if genetic differentiation coincides with the distributional range or morphological and plumage variation of described subspecies.

Greater spatial structuring and/or higher genetic diversity of western populations is noted from mtDNA studies of several North American birds that show east/west population differentiation (e.g. Kimura et al. 2002, Lovette et al. 2004, Klicka et al. 2011). This structure is most apparent for sedentary species or permanent resident populations of migratory species (e.g. Barrowclough et al. 2004, Spellman et al. 2007, Klicka et al. 2011, Manthey et al. 2011), but has also been described for migratory passerines (e.g. Kimura et al. 2002, Reugg and Smith 2002, Lovette et al. 2004), leading some authors to suggest that western populations might have undergone less severe reductions in effective population size during Pleistocene glaciation or might have occupied multiple, disjunct glacial refugia (Lovette et al. 2004, Klicka et al. 2011). Species distribution models for the American goldfinch identify a break in suitable breeding habitat during the LGM between northern and southern regions of the Pacific coast, indicating that genetic differentiation between western populations might stem from their occupation of isolated refugia during past glacial cycles.

The topographical complexity of the western mountain ranges and intervening xeric landscapes could also contribute to the current structuring observed among some western populations by limiting post-glaciation gene flow, although it remains unclear to what extent these features might influence differentiation in more vagile species (Calsbeek et al. 2003) and factors such as natal philopatry might play an equally important role (Manthay et al. 2011). We found that all breeding season individuals in western subclade 1 were sampled from California, whereas breeding season individuals in subclade 2 were all sampled from Washington state. However, in the nonbreeding season, some subclade 2 individuals were sampled from the same general region as subclade 1 (and both were recorded as *S. t. salicamans* in museum metadata), or from Southern California, which lies outside the currently described overwintering range for *S. t. jewetti*. It therefore remains unclear if the two subclades we identify here truly correspond to the described western subspecies, although it is possible that some *S. t. jewetti* individuals migrate further south than is currently believed and it should also be noted that some subspecies identifications are based on geographic location alone. These observations highlight the potential complexity of defining annual cycle connectivity in partially migratory species where populations, or even subspecies, might overlap in distributional range throughout portions of their annual cycle and further work is needed to assign individuals sampled during the nonbreeding period to their source breeding population. With sufficiently dense sampling, population assignment methods can successfully assign individuals to their breeding population even in cases with weak genetic differentiation among reference breeding populations and even when low-coverage whole-genome sequencing data is used (DeSaix et al. 2023a,b). Studying partially migratory species such as the American goldfinch, which has already demonstrated recent shifts in breeding distributional range (McCaslin and Heath 2020), could prove especially fruitful in understanding the evolutionary cost/benefit tradeoffs in migratory behavior associated with anthropogenic climate change and landscape-scale alterations to habitat.

In addition to historical demographic events and current gene flow, selection can contribute to current patterns of genetic differentiation by promoting local adaptation or assortative mating. We found only moderate per-site average Fst in pairwise comparison of the eastern and western populations (Fst= 0.0132) and very weak differentiation between the two western subclades (mean pairwise Fst= 0.0026), with no fixed SNP differences between populations in either comparison. There were also no Fst outlier SNPs when comparing the two western subclades. Outliers were identified between the eastern and western populations, but we found no enrichment for Gene Ontology terms and outlier SNPs almost exclusively fell outside of coding regions or caused only synonymous amino acid replacements. High-throughput sequencing has been instrumental in identifying genes underpinning migratory behavior and plumage patch coloration in several recent studies, where islands of genetic differentiation contained genes that were strongly associated with discrete phenotypic characters (Delmore et al. 2015, Mason and Taylor 2015, Toews et al. 2016). The underlying basis of traits exhibiting quantitative variation, as is more likely the case for clinal variation in body size and plumage tone or saturation in the American goldfinch, might be less easily determined (Harrison et al. 2012) and might also be attributable to environmental or dietary factors rather than to individual genetic variation. However, that many of the recently identified mutations involved in plumage coloration were localized to regulatory regions rather than to coding sequence (Funk and Taylor 2019) suggests that incorporating gene expression data could be a promising future avenue for research in the American goldfinch.

In conclusion, whole-genome resequencing provides clear evidence for population structuring in the American goldfinch despite relatively weak differentiation between populations and the potential confounding signatures of introgression or migration. Genome-wide analyses also indicate that past demographic events tied to historic climate cycles likely contributed to shaping present genetic diversity in this species even when accompanied by high long-term effective population size. Together, these results highlight the power of genome-level sequencing approaches to deepen our understanding of evolutionary processes in nonmodel wild species and to contribute to efforts assessing how contemporary factors might influence biodiversity. The fine-grained knowledge of how genetic variation is partitioned in widespread species that is afforded by genome-level analysis can highlight isolated subpopulations that might be of special concern due to small population size or unique local adaptations. Furthermore, responses of common and widespread species to anthropogenic climate change could carry outsized repercussions for rarer members of ecological assemblages through altered competition for resources or shifts in wildlife disease dynamics.

## Data availability statement

All sequencing data will be archived in the National Center for Biotechnology Information (NCBI). Accession numbers will be provided upon manuscript acceptance.

## Author contributions

Conceptualization: S.Y.W.S; sample collection: S.Y.W.S and E.S.K.P.; methodology: S.Y.W.S., E.S.K.P. and D.T.C.C.; investigation: E.S.K.P. and D.T.C.C.; analysis: A.C. and D.T.C.C.; visualization: A.C. and D.T.C.C.; writing—original draft: A.C.; writing—review and editing, all authors; supervision, project administration, and funding acquisition: S.Y.W.S.

## Acknowledgements

We thank the Field Museum, Louisiana State University Museum of Natural History, Museum of Comparative Zoology, University of California Museum of Vertebrate Zoology, and University of Washington Burke Museum for providing specimen loans. This project was supported by the startup fund (the University of Hong Kong) granted to S.Y.W.S. Computations were performed using research computing facilities offered by the Information Technology Services at the University of Hong Kong.

---

**Fig. S1.**
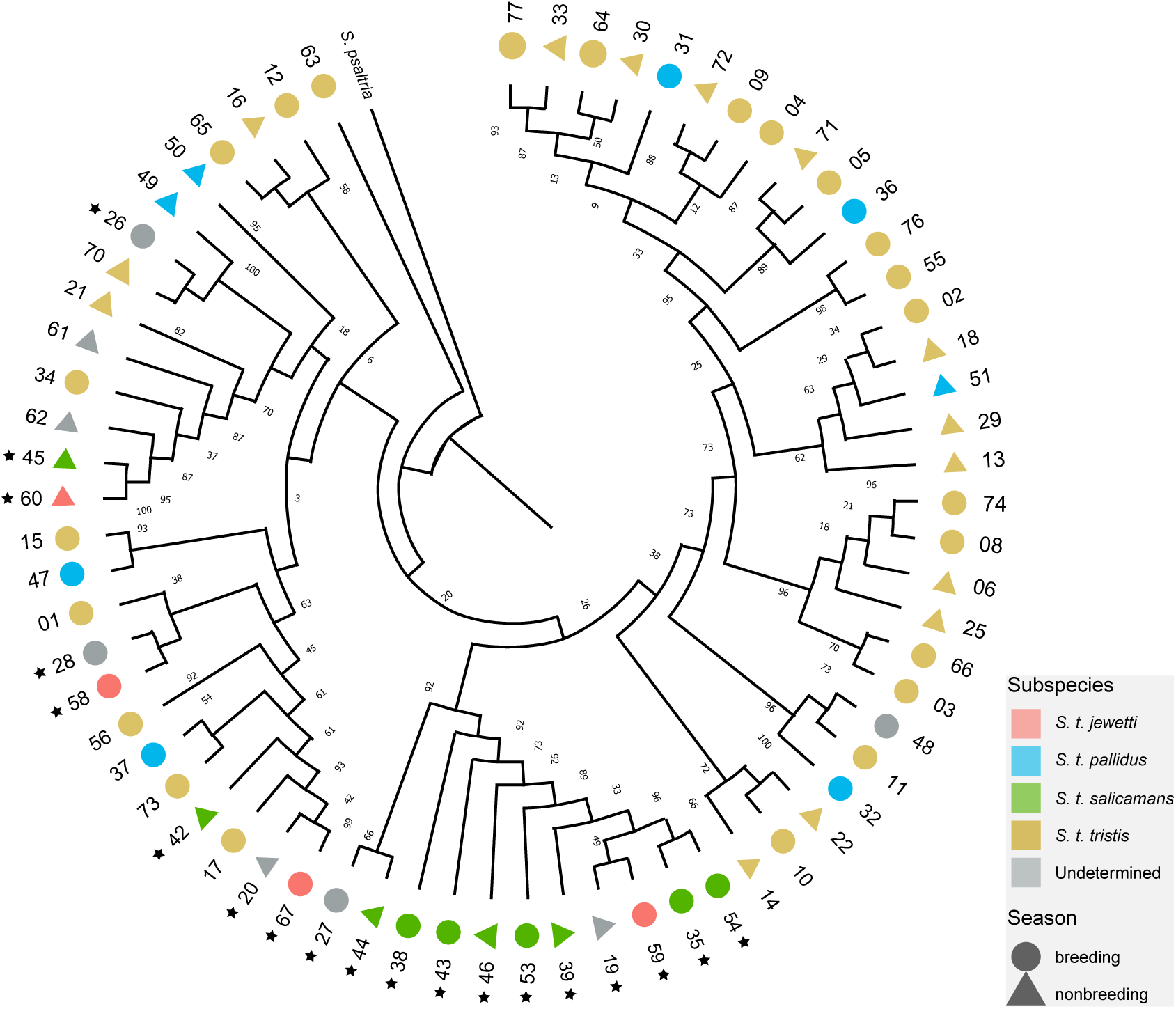
Mitochondrial DNA (mtDNA) phylogeny of *S. tristis* and *S. psaltria* (lesser goldfinch) outgroup. Branch lengths are uninformative and percentage support values from 500 bootstrap replicates are indicated. Individuals belonging to the western population from PCA and ADMIXTURE analyses (Fig. 2) are indicated with star symbols.

**Fig. S2.**
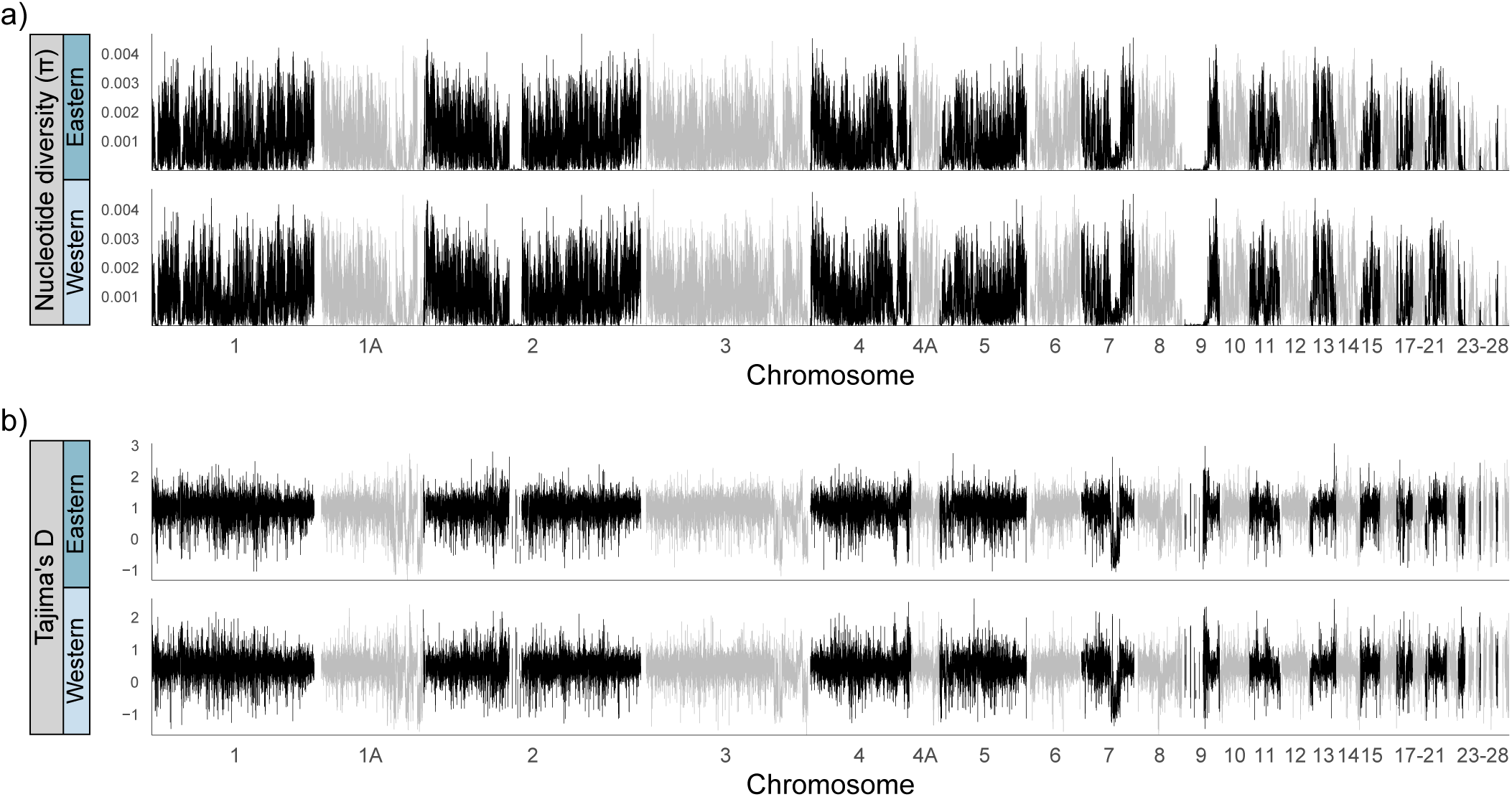
Genetic diversity and selection in eastern and western populations of *S. tristis*. Estimates of nucleotide diversity (π, part a), and Tajima’s D (b) are plotted along pseudochromosomes that were compiled by aligning *S. tristis* scaffolds to the zebra finch (*T. guttata*) reference assembly with SatsumaSynteny. Note that positional values within chromosomes correspond to the zebra finch coordinate system and data gaps where values fall to zero therefore represent regions of the zebra finch reference for which there was no aligned *S. tristis* sequence (for example, in chromosomes 2 & 9).

**Fig. S3.**
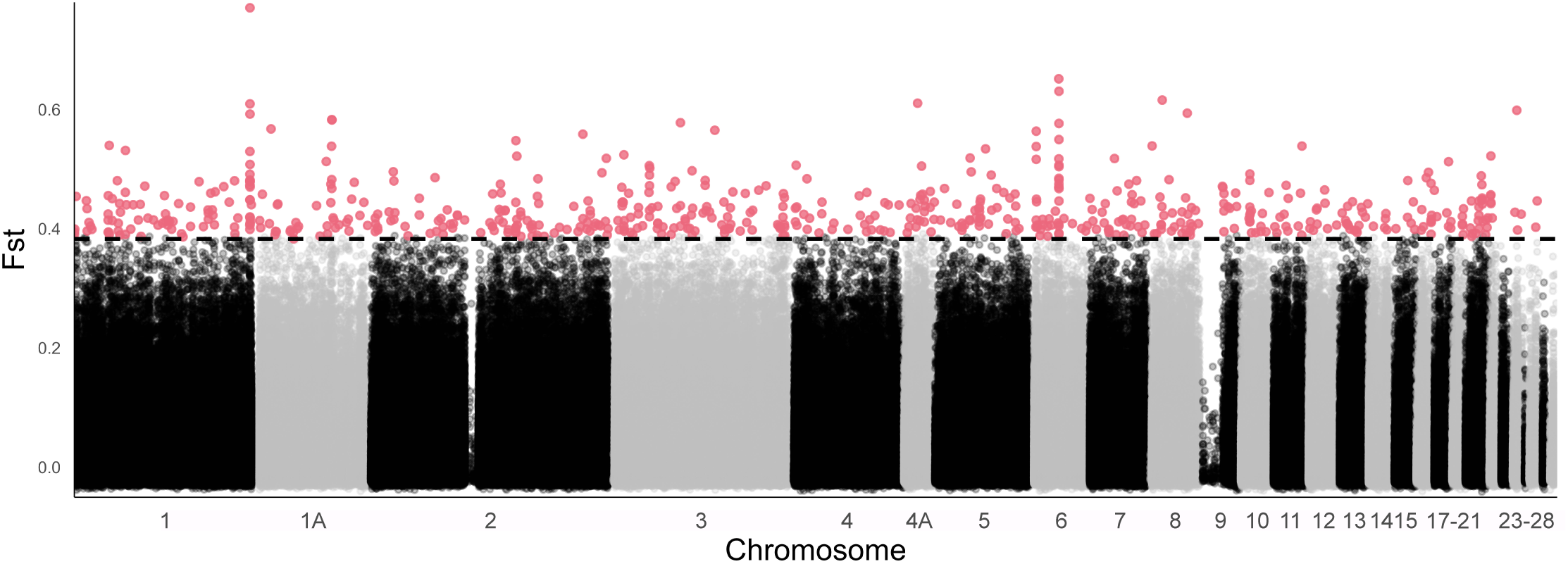
Fst outliers identified from comparing the eastern and western populations of *S. tristis* with OutFLANK. Per-site Fst values are plotted along pseudochromosomes that were compiled by aligning *S. tristis* scaffolds to the zebra finch (*T. guttata*) reference assembly with SatsumaSynteny. Fst values determined to be statistical outliers at a q-value threshold of 0.05 are shaded in pink and the significance cutoff is indicated with a dashed horizontal line. Note that positional values within chromosomes correspond to the zebra finch coordinate system and data gaps where values fall to zero therefore represent regions of the zebra finch reference for which there was no aligned *S. tristis* sequence (for example, in chromosomes 2 & 9).

**Table S1.** Summary of best candidate model selection for fastsimcoal2 simulations that estimated demographic parameters for the eastern and western populations of *Spinus tristis*. The best model chosen according to the minimum Akaike information criterion (AIC) is shown in bold, and ΔAIC of other models is shown relative to this best candidate model of ongoing migration between the eastern and western populations.

| Model | Max. Estimated Likelihood | Akaike Information Criterion (AIC) | ΔAIC |
| --- | --- | --- | --- |
| No migration | -53053513 | 244320467 | 2893415 |
| Migration in 2 different periods | -52426639 | 241433616 | 6564 |
| Early migration | -52966023 | 243917567 | 2490515 |
| **Ongoing migration** | **-52425215** | **241427052** | **0** |
| Recent migration | -52429839 | 241448347 | 21295 |
| Early migration followed by independent bottlenecks | -53054243 | 244323851 | 2896799 |
| Independent bottlenecks with no migration | -53055206 | 244328279 | 2901227 |
| Independent bottlenecks followed by migration | -52433641 | 241465868 | 38816 |

## References

1. Aguillon, S. M., Campagna, L., Harrison, R. G. & Lovette, I. J., 2018. A flicker of hope: genomic data distinguish Northern Flicker taxa despite low levels of divergence. The Auk, Volume 135, p. 748–766.

2. Alexander, D. H., Novembre, J. & Lange, K., 2009. Fast model-based estimation of ancestry in unrelated individuals. Genome Research, Volume 19, p. 1655–1664.

3. Andrews, S., 2010. FastQC: A quality control tool for high throughput sequence data. Available online at: http://www.bioinformatics.babraham.ac.uk/projects/fastqc/

4. Arnaiz-Villena, A. et al., 2012. Three different North American siskin/goldfinch evolutionary radiations (Genus *Carduelis*): pine siskin green morphs and European siskins in America. The Open Ornithology Journal, Volume 5, p. 73–81.

5. Arnaiz-Villena, A., Gomez-Prieto, P. & Ruiz-del-Valle, V., 2009. Phylogeography of finches and sparrows. In: Animal Genetics. L. J. Rechi, ed. Nova Science Publishers, Inc., p. 1–54.

6. Avise, J. C., 2000. Phylogeography: the history and formation of species. Harvard University Press, Cambridge, MA, U.S.A.

7. Avise, J. C. et al., 1987. Intraspecific phylogeography: the mitochondrial DNA bridge between population genetics and systematics. Annual Review of Ecology and Systematics, Volume 18, p. 489–522.

8. Avise, J. C. & Walker, D. E., 1998. Pleistocene phylogeographic effects on avian populations and the speciation process. Proceedings of the Royal Society B Biological Sciences, Volume 265, p. 457–463.

9. Barrowclough, G. F., Groth, J. G., Mertz, L. A. & Gutiérrez, R. J., 2004. Phylogeographic structure, gene flow and species status in blue grouse (*Dendragapus obscurus*). Molecular Ecology, Volume 13, p. 1911–1922.

10. Beckman, E. J. & Witt, C. C., 2015. Phylogeny and biogeography of the New World siskins and goldfinches: rapid, recent diversification in the Central Andes. Molecular Phylogenetics and Evolution, Volume 87, p. 28–45.

11. Beichman, A. C., Huerta-Sanchez, E. & Lohmueller, K. E., 2018. Using genomic data to infer historic population dynamics of nonmodel organisms. Annual Review of Ecology, Evolution, and Systematics, Volume 49, p. 433–456.

12. Bolger, A. M., Lohse, M. & Usadel, B., 2014. Trimmomatic: a flexible trimmer for Illumina sequence data. Bioinformatics, Volume 30, p. 2114–2120.

13. Brown, J. L., Bennett, J. R. & French, C. M., 2017. SDMtoolbox 2.0: the next generation Python-based GIS toolkit for landscape genetic, biogeographic and species distribution model analyses. PeerJ, Volume 5, p. e4095.

14. Brown, J. L. et al., 2018. PaleoClim, high spatial resolution paleoclimate surfaces for global land areas. Scientific Data, Volume 5, p. 180254.

15. Brunsfeld, S. J., Sullivan, J., Soltis, D. E. & Soltis, P. S., 2001. Comparative phylogeography of northwestern North America: A synthesis. In: Integrating Ecology and Evolution in a Spatial Context, J. Silvertown & J. Antonovics, Eds., Blackwell Science. p. 319–340.

16. Burbrink, F. T. et al., 2016. Asynchronous demographic responses to Pleistocene climate change in Eastern Nearctic vertebrates. Ecology Letters, Volume 19, p. 1457–1467.

17. Burri, R. et al., 2015. Linked selection and recombination rate variation drive the evolution of the genomic landscape of differentiation across the speciation continuum of Ficedula flycatchers. Genome Research, Volume 25, p. 1656–1665.

18. Bushnell, B. (2014). BBMap: A fast, accurate, splice-aware aligner. Lawrence Berkeley National Laboratory. LBNL Report #: LBNL-7065E. Retrieved from: https://escholarship.org/uc/item/1h3515gn

19. Calsbeek, R., Thompson, J. N. & Richardson, J. E., 2003. Patterns of molecular evolution and diversification in a biodiversity hotspot: the California Floristic Province. Molecular Ecology, Volume 12, p. 1021–1029.

20. Capella-Gutiérrez, S., Silla-Martínez, J. M. & Gabaldón, T., 2009. trimAl: a tool for automated alignment trimming in large-scale phylogenetic analyses. Bioinformatics, Volume 25, p. 1972–1973.

21. Chang, C. C. et al., 2015. Second-generation PLINK: rising to the challenge of larger and richer datasets. GigaScience, Volume 4, p. 7.

22. Chen, S., Zhou, Y., Chen, Y. & Gu, J., 2018. fastp: an ultra-fast all-in-one FASTQ preprocessor. Bioinformatics, Volume 34, p. i884–i890.

23. Cheviron, Z. A. & Swanson, D. L., 2017. Comparative transcriptomics of seasonal phenotypic flexibility in two North American songbirds. Integrative and Comparative Biology, Volume 57, p. 1040–1054.

24. Colbeck, G. J. et al., 2008. Phylogeography of a widespread North American migratory songbird (*Setophaga ruticilla*). The Journal of Heredity, Volume 99, p. 453–463.

25. Constable, H. et al., 2010. VertNet: a new model for biodiversity data sharing. PLoS Biology, Volume 8, p. e1000309.

26. Cristofari, R. et al., 2018. Climate-driven range shifts of the king penguin in a fragmented ecosystem. Nature Climate Change, Volume 8, p. 245–251.

27. Danecek, P. et al., 2011. The variant call format and VCFtools. Bioinformatics, Volume 27, p. 2156–2158.

28. Danecek, P. et al., 2021. Twelve years of SAMtools and BCFtools. GigaScience, Volume 10, p. 1–4.

29. Delmore, K. E. et al., 2015. Genomic analysis of a migratory divide reveals candidate genes for migration and implicates selective sweeps in generating islands of differentiation. Molecular Ecology, Volume 24, p. 1873–1888.

30. DeSaix, M. G. et al., 2023a. Low-coverage whole genome sequencing for highly accurate population assignment: Mapping migratory connectivity in the American Redstart (*Setophaga ruticilla*). Molecular Ecology, Volume 32, p. 5528–5540.

31. DeSaix, M. G., Rodriguez, M. D., Ruegg, K. C., & Anderson, E. C., 2023b. Population assignment from genotype likelihoods for low-coverage whole-genome sequencing data. Authorea, 1–47. Retrieved from: 10.22541/au.168569102.27840692/v1

32. Dierickx, E.G., Sin, S.Y.W., van Veelen, H.P.J., Brooke, M.D.L., Liu, Y., Edwards, S.V. & Martin, S.H., 2020. Genetic diversity, demographic history and neo-sex chromosomes in the Critically Endangered Raso lark. Proceedings of the Royal Society B, Volume 287, p.20192613.

33. Dutoit, L. et al., 2017. Covariation in levels of nucleotide diversity in homologous regions of the avian genome long after completion of lineage sorting. Proceedings of the Royal Society B Biological Sciences, Volume 284, p. 20162756–20162756.

34. eBird. 2021. eBird: an online database of bird distribution and abundance [web application]. eBird, Cornell Lab of Ornithology, Ithaca, New York. Available: http://www.ebird.org. (Data Version: 2019, Accessed: June 11, 2020).

35. Edwards, S. V., Robin, V. V., Ferrand, N. & Moritz, C., 2022. The evolution of comparative phylogeography: putting the geography (and more) into comparative population genomics. Genome Biology and Evolution, Volume 14, p. evab176.

36. Evanno, G., Regnaut, S. & Goudet, J., 2005. Detecting the number of clusters of individuals using the software structure: a simulation study. Molecular Ecology, Volume 14, p. 2611–2620.

37. Excoffier, L. et al., 2013. Robust demographic inference from genomic and SNP data. PLoS Genetics, Volume 9, p. e1003905.

38. Excoffier, L. et al., 2021. fastsimcoal2: demographic inference under complex evolutionary scenarios. Bioinformatics, Volume 37, p. 4882–4885.

39. Francis, R. M., 2017. pophelper: an R package and web app to analyse and visualize population structure. Molecular Ecology Resources, Volume 17, p. 27–32.

40. Funk, E. R. & Taylor, S. A., 2019. High-throughput sequencing is revealing genetic associations with avian plumage color. The Auk, Volume 136, p. 1–7.

41. Gagnaire, P., 2020. Comparative genomics approach to evolutionary process connectivity. Evolutionary Applications, Volume 13, p. 1320–1334.

42. García-Roselló, E. et al., 2013. ModestR: a software tool for managing and analyzing species distribution map databases. Ecography, Volume 36, p. 1202–1207.

43. Grabherr, M. G. et al., 2010. Genome-wide synteny through highly sensitive sequence alignment: Satsuma. Bioinformatics, Volume 26, p. 1145–1151.

44. Grinnell, J., 1897. New race of *Spinus tristis* from the Pacific Coast. The Auk, Volume 14, p. 397–399.

45. Gutenkunst, R. N., Hernandez, R. D., Williamson, S. H. & Bustamante, C. D., 2009. Inferring the joint demographic history of multiple populations from multidimensional SNP frequency data. PLoS Genetics, Volume 5, p. e1000695.

46. Hahn, C., Bachmann, L. & Chevreux, B., 2013. Reconstructing mitochondrial genomes directly from genomic next-generation sequencing reads–a baiting and iterative mapping approach. Nucleic Acids Research, Volume 41, p. e129.

47. Hanghøj, K. et al., 2019. Fast and accurate relatedness estimation from high-throughput sequencing data in the presence of inbreeding. GigaScience, Volume 8, p. 1–8.

48. Harrison, P. W., Wright, A. E. & Mank, J. E., 2012. The evolution of gene expression and the transcriptome–phenotype relationship. Seminars in Cell & Developmental Biology, Volume 23, p. 222–229.

49. Hewitt, G., 2000. The genetic legacy of the Quaternary ice ages. Nature, Volume 405, p. 907– 913.

50. Hewitt, G. M., 2004. The structure of biodiversity - insights from molecular phylogeography. Frontiers in Zoology, Volume 1, p. 4.

51. Huerta-Cepas, J., Serra, F. & Bork, P., 2016. ETE 3: reconstruction, analysis, and visualization of phylogenomic data. Molecular Biology and Evolution, Volume 33, p. 1635–1638.

52. Huynh, S., Cloutier, A., Chen, G., Chan, D.T.C., Lam, D.K., Huyvaert, K.P., Sato, F., Edwards, S.V. & Sin, S.Y.W., 2023. Whole-genome analyses reveal past population fluctuations and low genetic diversities of the North Pacific albatrosses. Molecular Biology and Evolution, Volume 40, p.msad155.

53. IUCN. 2016. The IUCN Red List of Threatened Species. Version 2016-3. Available at: www.iucnredlist.org. Accessed on December 10, 2023.

54. Kalyaanamoorthy, S. et al., 2017. ModelFinder: fast model selection for accurate phylogenetic estimates. Nature Methods, Volume 14, p. 587–589.

55. Katoh, K. & Standley, D. M., 2013. MAFFT multiple sequence alignment software version 7: improvements in performance and usability. Molecular Biology and Evolution, Volume 30, p. 772–780.

56. Kelly, R. J., Murphy, T. G., Tarvin, K. A. & Burness, G., 2012. Carotenoid-based ornaments of female and male American Goldfinches (*Spinus tristis*) show sex-specific correlations with immune function and metabolic rate. Physiological and Biochemical Zoology, Volume 85, p. 348–363.

57. Kimura, M. et al., 2002. Phylogeographical approaches to assessing demographic connectivity between breeding and overwintering regions in a Nearctic−Neotropical warbler (*Wilsonia pusilla*). Molecular Ecology, Volume 11, p. 1605–1616.

58. Klicka, J. et al., 2011. A phylogeographic and population genetic analysis of a widespread, sedentary North American bird: the Hairy Woodpecker (*Picoides villosus*). The Auk, Volume 128, p. 346–362.

59. Kopelman, N. M. et al., 2015. CLUMPAK: a program for identifying clustering modes and packaging population structure inferences across K. Molecular Ecology Resources, Volume 15, p. 1179–1191.

60. Li, H. & Durbin, R., 2009. Fast and accurate short read alignment with Burrows–Wheeler transform. Bioinformatics, Volume 25, p. 1754–1760.

61. Li, H. & Durbin, R., 2011. Inference of human population history from individual whole-genome sequences. Nature, Volume 475, p. 493–496.

62. Li, H. et al., 2009. The sequence alignment/map format and SAMtools. Bioinformatics, Volume 25, p. 2078–2079.

63. Li, S. et al., 2014. Genomic signatures of near-extinction and rebirth of the crested ibis and other endangered bird species. Genome Biology, Volume 15, p. 557–557.

64. Linnaeus, C., 1758. Systema Naturae per regna tria naturae, secundum classes, ordines, genera, species, cum characteribus, differentiis, synonymis, locis. Editio decima, reformata [10th revised edition]

65. Liu, X. & Fu, Y.-X., 2015. Exploring population size changes using SNP frequency spectra. Nature Genetics, Volume 47, p. 555–559.

66. Liu, X. & Fu, Y.-X., 2020. Stairway Plot 2: demographic history inference with folded SNP frequency spectra. Genome Biology, Volume 21, p. 280.

67. Lovette, I. J., 2005. Glacial cycles and the tempo of avian speciation. Trends in Ecology & Evolution, Volume 20, p. 57–59.

68. Lovette, I. J., Clegg, S. M. & Smith, T. B., 2004. Limited utility of mtDNA markers for determining connectivity among breeding and overwintering locations in three neotropical migrant birds. Conservation Biology, Volume 18, p. 156–166.

69. Lyman, R. A. & Edwards, C. E., 2022. Revisiting the comparative phylogeography of unglaciated eastern North America: 15 years of patterns and progress. Ecology and Evolution, Volume 12, p. e8827.

70. Mable, B. K., 2019. Conservation of adaptive potential and functional diversity: integrating old and new approaches. Conservation Genetics, Volume 20, p. 89–100.

71. Manthey, J. D., Klicka, J. & Spellman, G. M., 2011. Cryptic diversity in a widespread North American songbird: phylogeography of the Brown Creeper (*Certhia americana*). Molecular Phylogenetics and Evolution, Volume 58, p. 502–512.

72. Manthey, J. D., Klicka, J. & Spellman, G. M., 2021. The genomic signature of allopatric speciation in a songbird is shaped by genome architecture (Aves: *Certhia americana*). Genome Biology and Evolution, Volume 13, p. evab120.

73. Mason, N. A. & Taylor, S. A., 2015. Differentially expressed genes match bill morphology and plumage despite largely undifferentiated genomes in a Holarctic songbird. Molecular Ecology, Volume 24, p. 3009–3025.

74. McCaslin, H. M. & Heath, J. A., 2020. Patterns and mechanisms of heterogeneous breeding distribution shifts of North American migratory birds. Journal of Avian Biology, Volume 2020, p. e02237.

75. McGraw, K. J. & Middleton, A. L., 2020. American Goldfinch (*Spinus tristis*), version 1.0. In: Birds of the World, P. G. Rodewald, Editor. Cornell Lab of Ornithology, Ithaca, NY, USA. https://doi-org.proxy.lib.sfu.ca/10.2173/bow.amegfi.01

76. Mearns, E. A., 1890. Descriptions of a new species and three new subspecies of birds from Arizona. The Auk, Volume 7, p. 243–251.

77. Middleton, A. L. A., 1978. The annual cycle of the American Goldfinch. The Condor, Volume 80, p. 401–406.

78. Mi, H. et al., 2019. Protocol update for large-scale genome and gene function analysis with the PANTHER classification system (v.14.0). Nature Protocols, Volume 14, p. 703–721.

79. Milà, B., Girman, D. J., Kimura, M. & Smith, T. B., 2000. Genetic evidence for the effect of a postglacial population expansion on the phylogeography of a North American songbird. Proceedings of the Royal Society B Biological Sciences, Volume 267, p. 1033–1040.

80. Milot, E., Gibbs, H. L. & Hobson, K. A., 2000. Phylogeography and genetic structure of northern populations of the yellow warbler (*Dendroica petechia*). Molecular Ecology, Volume 9, p. 667–681.

81. Minh, B. Q. et al., 2020. IQ-TREE 2: new models and efficient methods for phylogenetic inference in the genomic era. Molecular Biology and Evolution, Volume 37, p. 1530–1534.

82. Miraldo, A. et al., 2016. An Anthropocene map of genetic diversity. Science, Volume 353, p. 1532–1535.

83. Moreira, L. R., Klicka, J. & Smith, B. T., 2023. Demography and linked selection interact to shape the genomic landscape of codistributed woodpeckers during the Ice Age. Molecular Ecology, Volume 32, p. 1739–1759.

84. Nadachowska-Brzyska, K. et al., 2015. Temporal dynamics of avian populations during Pleistocene revealed by whole-genome sequences. Current Biology, Volume 25, p. 1375–1380.

85. Partners in Flight. 2020. Population Estimates Database, version 3.1. Available at http://pif.birdconservancy.org/PopEstimates. Accessed on December 10, 2023.

86. Pedersen, B. S. & Quinlan, A. R., 2018. Mosdepth: quick coverage calculation for genomes and exomes. Bioinformatics, Volume 34, p. 867–868.

87. Phillips, S. J. et al., 2017. Opening the black box: an open-source release of Maxent. Ecography, Volume 40, p. 887–893.

88. Prescott, D. R. C. & Middleton, A. L. A., 1990. Age and sex differences in winter distribution of American Goldfinches in eastern North America. Ornis Scandinavica, Volume 21, p. 99– 104.

89. QGIS Development Team, 2021. QGIS Geographic Information System. Retrieved from: http://qgis.org

90. Quinlan, A. R. & Hall, I. M., 2010. BEDTools: a flexible suite of utilities for comparing genomic features. Bioinformatics, Volume 26, p. 841–842.

91. R Core Team, 2021. R: A language and environment for statistical computing. R Foundation for Statistical Computing, Vienna, Austria. Available at: https://www.R-project.org/.

92. Robinson, J. et al., 2016. Genomic flatlining in the endangered Island Fox. Current Biology, Volume 26, p. 1183–1189.

93. Ruegg, K. C. & Smith, T. B., 2002. Not as the crow flies: a historical explanation for circuitous migration in Swainson’s thrush (*Catharus ustulatus*). Proceedings of the Royal Society B Biological Sciences, Volume 269, p. 1375–1381.

94. Sin, S.Y.W., Hoover, B.A., Nevitt, G.A. & Edwards, S.V., 2021. Demographic history, not mating system, explains signatures of inbreeding and inbreeding depression in a large outbred population. The American Naturalist, Volume 197, p. 658–676.

95. Smith, B. T., Gehara, M. & Harvey, M. G., 2021. The demography of extinction in eastern North American birds. Proceedings of the Royal Society B Biological sciences, Volume 288, p. 20201945.

96. Smith, B. T. et al., 2017. A latitudinal phylogeographic diversity gradient in birds. PLoS Biology, Volume 15, p. e2001073.

97. Soltis, D. E. et al., 2006. Comparative phylogeography of unglaciated eastern North America. Molecular Ecology, Volume 15, p. 4261–4293.

98. Spellman, G. M., Riddle, B. & Klicka, J., 2007. Phylogeography of the mountain chickadee (*Poecile gambeli*): diversification, introgression, and expansion in response to Quaternary climate change. Molecular Ecology, Volume 16, p. 1055–1068.

99. Stamatakis, A., 2014. RAxML version 8: a tool for phylogenetic analysis and post-analysis of large phylogenies. Bioinformatics, Volume 30, p. 1312–1313.

100. Sughrue, K. M., Brittingham, M. C. & French, J. B., 2008. Endocrine effects of the herbicide linuron on the American Goldfinch (*Carduelis tristis*). The Auk, Volume 125, p. 411–419.

101. Tacutu, R. et al., 2018. Human ageing genomic resources: new and updated databases. Nucleic Acids Research, Volume 46, p. D1083–D1090.

102. Theodoridis, S. et al., 2020. Evolutionary history and past climate change shape the distribution of genetic diversity in terrestrial mammals. Nature Communications, Volume 11, p. 2557.

103. Thomas, P. D. et al., 2022. PANTHER: Making genome-scale phylogenetics accessible to all. Protein Science, Volume 31, p. 8–22.

104. Toews, D. L. et al., 2016. Plumage genes and little else distinguish the genomes of hybridizing warblers. Current Biology, Volume 26, p. 2313–2318.

105. Van der Auwera, G. A. et al., 2013. From FastQ data to high-confidence variant calls: the Genome Analysis Toolkit best practices pipeline. Current Protocols in Bioinformatics, Volume 43, p. 11.10.1–11.10.33.

106. Van der Auwera, G. A. & O’Connor, B. D., 2020. Genomics in the cloud: using Docker, GATK, and WDL in Terra. First edition. O’Reilly Media.

107. van Rossem, A. J., 1943. Description of a race of goldfinch from the Pacific Northwest. The Condor, Volume 45, p. 158–159.

108. Weisenfeld, N. I. et al., 2017. Direct determination of diploid genome sequences. Genome Research, Volume 27, p. 757–767.

109. Wickham, H., 2016. ggplot2: elegant graphics for data analysis. 2nd ed. 2016 ed. Cham: Springer International Publishing.

110. Wolf, J. B. W., Lindell, J. & Backström, N., 2010. Speciation genetics: current status and evolving approaches. Philosophical Transactions of the Royal Society of London Series B Biological sciences, Volume 365, p. 1717–1733.

111. Yang, Z., 2014. Molecular Evolution: A Statistical Approach. Oxford University Press, Oxford, U.K.

112. Zink, R. M., 1996. Comparative phylogeography in North American birds. Evolution, Volume 50, p. 308–317.

